# Role of a childhood cancer-linked BRIP1/FANCJ germline variant in genomic instability and cancer cell vulnerability

**DOI:** 10.64898/2026.03.24.714005

**Authors:** Sahar Karbassi, Teodora Nikolova, Daniela Pfeiffer, Swaroopa Nakkeeran, Roopesh Anand, Pierre-Olivier Frappart, Michaela Kuhlen, Thomas G Hofmann

**Author notes:** Correspondence should be addressed to T.G.H.

## Abstract

Childhood cancer is frequently associated with inherited pathogenic variants of cancer predisposition genes. Using Whole-Exome Sequencing, we identified an inherited, monoallelic pediatric cancer-linked germline variant of DNA helicase FANCJ/BRIP1, BRIP1^R162Q^ of unclear clinical significance. Intriguingly, *in vitro* helicase assays demonstrated that BRIP1^R162Q^ encodes a hyperactive DNA helicase. In cells, stable expression of BRIP1^R162Q^ in BRIP1-proficient cells selectively sensitized to ectopic DNA replication stress. Moreover, BRIP1^R162Q^ expressing cells showed chronic, steady-state activation of DNA replication stress indicated by decreased fork speed, fork stalling and impaired fork symmetry. Moreover, BRIP1^R162Q^ expression triggered genomic instability indicated by elevated γH2AX foci numbers and chromosomal aberrations, suggesting BRIP1^R162Q^ as a variant driving genomic instability. Mechanistically, BRIP1^R162Q^ mislocalizes upon replication stress, and leads to accumulation of DNA secondary G-quadruplex (G4) structures and R-loops. RNaseH1 expression, which resolves R-loops, also reduced G4 levels and relieved BRIP1^R162Q^-induced replication stress. Finally, BRIP1^R162Q^ introduces exploitable vulnerabilities for targeted therapies using G4-ligand Pyridostatin and DNA damage kinase ATR and DNA-PK inhibition. Our findings propose a mechanistic framework of how a childhood cancer-linked monoallic, hypermorphic BRIP1 germline variant participates in cancer development, and suggest potential therapeutic treatment strategies.

## Introduction

While lifestyle-dependent carcinogenic exposures and infectious agents are considered major contributors to adult-onset cancers, childhood cancer is frequently associated with pathogenic inherited small nucleotide variants (SNVs) in cancer predisposition genes. Well-established examples include TP53, RB1, NF1, and APC, whose germline variants are unambiguously linked to pediatric cancer predisposition syndromes^1^. In addition, large-scale whole exome sequencing (WES) studies of pediatric cancer cohorts have identified germline variants in Fanconi Anemia (FA)/breast cancer-associated (BRCA) pathway genes, which play a fundamental role in interstrand DNA crosslink repair (ICL) and homologous recombination (HR)^1^. While biallelic variants in FA/BRCA pathway genes such as BRCA2 (FANCD1), PALB2 (FANCN), and BRIP1 (FANCJ) cause Fanconi anemia with severe childhood cancer predisposition, the significance of heterozygous variants in these genes, canonically associated with adult-onset breast, ovarian, and pancreatic cancer, remains less well understood in the pediatric context^1, 2^. Notably, although such variants have been recurrently identified in pediatric WES studies, functional evidence supporting their causal role in childhood tumorigenesis is largely lacking, and a second somatic hit in the tumor has rarely been demonstrated^1, 3^. Next-generation sequencing of parent–child trios provides a powerful and cost-effective method to identify candidate pathogenic SNVs in cancer predisposition genes, yet functional validation remains essential to establish causality^4, 5^. Here, we functionally characterize a childhood cancer-linked germline variant in BRIP1/FANCJ and investigate its role in genomic instability and cancer cell vulnerability BRIP1/FANCJ/BACH1 is a DNA helicase with essential functions in regulating genomic stability by participating in DNA interstrand crosslink repair, Homogous recombination (HR) repair, G-quadruplex (G4) recognition and resolution, and DNA replication stress ^6–13^. Interestingly, BRIP1 germline variants have been identified in women with early-onset breast cancer, and these variants showed a compromised DNA helicase activity, suggesting that BRIP1 variants with compromised helicase activity participate in cancer development ^14, 15^. Similar, rare BRIP1 germline variants with reduced helicase activity have been identified in hereditary breast and ovarian cancer syndrome families^16, 17^.

Important insights into the role of BRIP1 in maintaining genome stability came from findings implicating the DNA helicase in recognizing and remodeling the genomic G4 landscape^11, 13^. Although the human genome harbors more than 700,000 potential G4-forming sequences^18^, only a minor fraction of them form G4 secondary structures in a natural chromatin context in the cell^19^, and moreover, different cell types form specific G4 subsets^20, 21^. In addition, G4’s can spontaneously form during DNA replication^22^ or transcription^23^, and unresolved G4s can trigger DNA damage^24^ linked to genomic deletions^25^ and rearreangements, and consequently G4’s have been linked to cancer^26^ and aging^27^. Deregulation of the genomic G4 landscape leads to altered gene expression profiles^20^ and interferes with normal differentiation of human embryonic stem cells^28^. G4s are frequently found in the vicinity of R-loops, which are three-stranded structures consisting of a RNA-DNA hybrid and a displaced ssDNA strand^29^. R-loops have regulatory functions in transcription, telomere stability and DNA repair^29^. However, unscheduled, persistent R-loop accumulation results in DNA replication stress, DNA double-strand breaks and genomic instability^29^. Interestingly, BRIP1//FANCJ plays a fundamental role in recognizing^11^ and resolving G4s and so-called G-loops^13^. G-loops consist of a G4 that forms on the displaced ssDNA strand a RNA:DNA hybrid termed R-loop, which stabilizes the G4 structure^13^. By regulating G-loop disassembly, BRIP1/FANCJ shapes the genomic G4 landscape, and deregulated BRIP1 function results in G4 and R-loop accumulation, leading to genomic instability.^13^

Since it is difficult to unequivocally deduce from *in silico* predictions whether identified SNVs in a cancer predisposition gene are bystanders (due to increased genomic instability) or are drivers of carcinogenesis, functional analysis is required to determine the clinical significance.

Here we provide a proof-of-principle study to evaluate of the biological effects of a rare BRIP1^R162Q^ germline variant of uncertain clinical significance linked to childhood cancer development. Our findings indicate that BRIP1^R162Q^ encodes a hyperactive DNA helicase, which triggers steady-state, chronic DNA replication stress and genomic instability. Mechanistically, we found that BRIP1^R162Q^ variant expression leads to an increase in DNA secondary G4-quadruplex structures and R-loops, RNA:DNA hybrids linked to DNA replication stress. Finally, we provide evidence that the BRIP1^R162Q^ variant introduces vulnerabilities in cancer cells which can be exploited for targeted cancer therapy.

## Results

### Identification of the rare BRIP1^R162Q^ germline variant in a child with cancer

To identify potentially pathogenic germline variants in DNA repair and DNA damage response factors linked to childhood cancer formation, we performed WES in parent-child trios as reported previously^30, 31^. We focussed on a 11 year-old girl with metastatic osteosarcoma, with a remarable family tree with cancer history in different generations (**Fig. 1a**). Functional testing performed with patient-derived peripheral blood mononuclear cells revealed no hypersensitivity to the DNA crosslinker mitomycin (MMC) (not shown)., excluding a classical Fanconi anemia phenotype (caused by biallelic FANC gene defects), which are linked to characteristic DNA-crosslinker hypersensitivity. WES data analyses by our bioinformatics pipeline identified a potentially pathogenic monoallelic single nucleotide variant of BRIP1/FANCJ (c.485G>A; p.R162Q), BRIP1^R162Q^, and a monoallelic variant of the DNA damage kinase HIPK2^32, 33^ (c.1804A>C;p.T602P) (**Fig. 1b**). While the BRIP1 variant has been predicted *in silico* as “probably damaging deleterious”, the HIPK2 variant was predicted to be tolerated and non-pathogenic. Interestingly, the rare BRIP1^R162Q^ variant has been reported previously as a “damaging” monoallelic hereditary variant in a patient with early-onset head and neck squamous cell carcinoma^34^. Due to lack of functional data, the BRIP1^R162Q^ variant is currently of uncertain clinical significance. Therefore, we aimed to investigate its functional impact.

**Figure 1.**
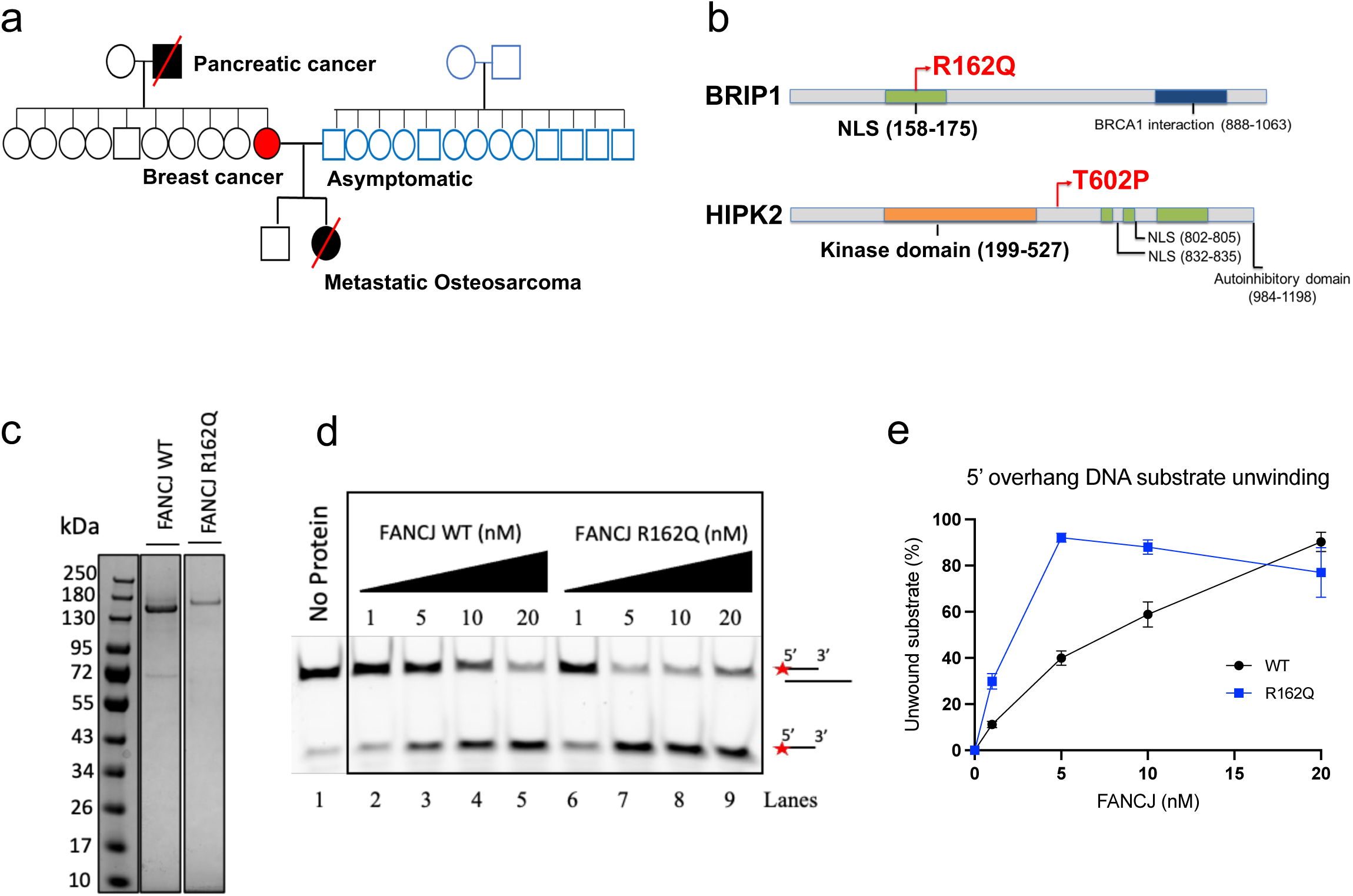
Whole-exome sequencing (WES) of a parent–child trio identifies a rare, hyper-active FANCJ/BRIP1^R162Q^ variant linked to pediatric cancer. **a,** Three-generation pedigree of an 11-year-old pediatric patient diagnosed with cancer investigated in this study. **b,** Whole-exome sequencing (WES) of peripheral-blood-derived DNA from the patient and both parents identified a novel missense variant in *BRIP1* (c.485G>A, p.Arg162Gln) located in the nuclear localization signal (NLS), inherited from the mother, and a missense variant in *HIPK2* (c.1804A>C, p.Thr602Pro), inherited from the father. **c,** Coomassie-stained SDS-PAGE of purified recombinant BRIP1/FANCJ WT and BRIP/FANCJ^R162Q^ proteins. **d,** FAM-labeled 5’ overhang DNA substrate unwinding assay using increasing concentrations of BRIP1 WT or R162Q (1-20 nM) as indicated. Upper signal indicates the duplex substrate, the lower one the unwound FAM-labelled oligonucleotide. A representative assay is shown **e,** Quantification of DNA unwinding assays (n=3) shown as percentage of unwound substrate (mean ± SEM).

### The cancer-linked BRIP1^R162Q^ variant encodes a hyperactive DNA helicase protein

BRIP1 is a DNA helicase enzyme exerting 5’-3’-unwinding activity, which can be biochemically analysed *in vitro* ^11, 12, 35, 36^. To determine whether the BRIP1^R162Q^ variant shows an altered helicase activity, we purified recombinant wild-type BRIP1 and the BRIP1^R162Q^ variant (**Fig.1c**) and tested their DNA unwinding activity in parallel using 5’-overhang substrate. Intriguingly, DNA unwinding analysis demonstrated that BRIP1^R162Q^ shows an about 3-fold increased DNA helicase unwinding activity compared to the wild-type BRIP1 protein (**Fig.1d, e**). In addition, a comparable hyperactivity of the BRIP1^R162Q^ variant was detected when an oligo forming a G4 DNA secondary structure was in G4-unwinding unwinding assays (**Supplementary Fig. 1 a,b**), strengthening our conclusion that the BRIP1^R162Q^ variant represents a hyperactive DNA helicase. In sum, our *in vitro* results indicate that the childhood cancer-linked BRIP1^R162Q^ variant is hypermorphic and shows hyperactive DNA helicase activity.

### BRIP1^R162Q^ increases cellular sensitivity to DNA replication stress

As the BRIP1^R162Q^ variant encodes a hyperactive DNA helicase *in vitro*, we next investigated the functional consequences of its expression in human cells. As experimental system we used human HCT116 colon cancer cells as they are nearly diploid, express wild-type BRIP1 and wild-type p53, an important regulator of the DNA damage response, and provide a well-established cell model to study cellular responses to DNA damage^37^. Since we identified BRIP1^R162Q^ as a monoallelic childhoood cancer-linked germline variant – indicating that it is expressed along with one BRIP1 wild-type allele - we tried to mimic the setting of co-expression of BRIP1 wild-type and the BRIP1^R162Q^ variant of the childhood cancer patient. As strategy we used lentiviral transduction of HA-Myc-BRIP1 wild-type or the HA-Myc-BRIP1^R162Q^ variant, and the empty vector as control, into BRIP1-proficient HCT116 cells to reveal cell pools with stable expression (**Fig. 2a**). The obtained cell pools were analysed by immunoblotting, which confirmed comparable expression of the HA-Myc-tagged BRIP1 proteins, along with the endogenously expressed wild-type BRIP1 protein (**Fig. 2c,d**). These stable cell pools were used throughout our entire study. In addition, we also generated BRIP1 knock-out (KO) HCT116 cells using CRISPR/Cas9-mediated gene editing (**Supplementary Fig. 2a**), and confirmed the loss of BRIP1 expression using DNA sequencing of the BRIP1 locus (**Supplementary Fig. 2b**) and immunoblotting (**Supplementary Fig. 2c**). Since BRIP1 is known to participate in ICL and HR repair^36, 38, 39^, we next challenged the BRIP1 WT, BRIP1^R162Q^ and BRIP1 KO cells with the DNA crosslinking agent Mitomycin C (MMC) and ionizing radiation (IR) to determine if the BRIP1^R162Q^ variant sensitizes to these DNA damaging treatment. Although colony formation assays revealed hypersensitivity of BRIP1 KO cells to both treatments, no difference between wild-type BRIP1 and the BRIP1^R162Q^ variant expressing cells was found in the sensitivity to MMC (**Fig. 2d,e**) and IR damage (**Fig. 2f,g**). However, when cells were exposed to DNA replication stress using hydroxy urea (HU), BRIP1^R162Q^ expressing cells showed increased sensitivity when compared to the wild-type counterparts (**Fig. 2h,i**). Thus, our results indicate that expression of BRIP1^R162Q^ is linked to a specific defect in the DNA replication stress response.

**Figure 2.**
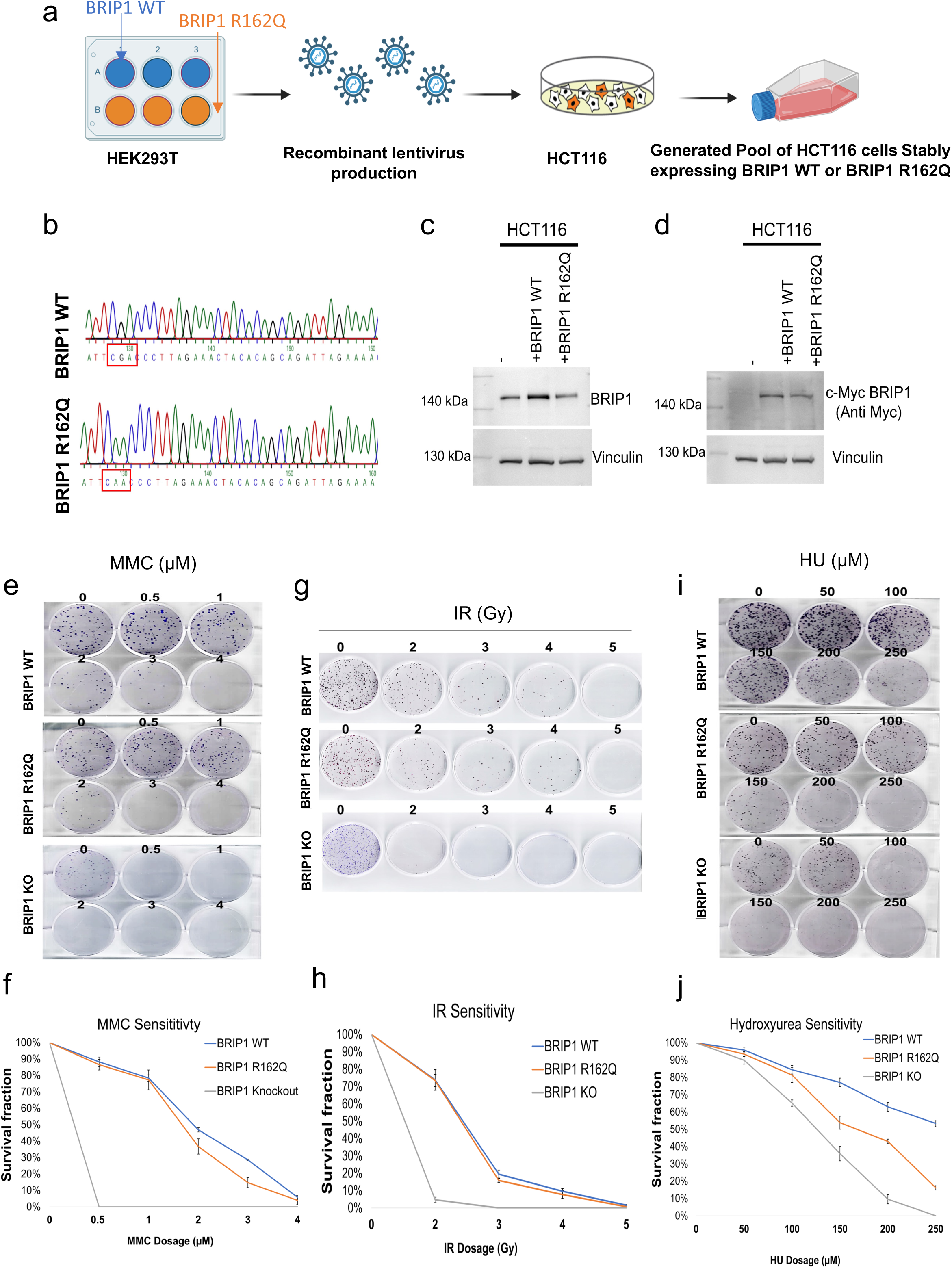
Pediatric cancer–linked BRIP1^R162Q^ variant selectively increases sensitivity to DNA replication stress. **a,** Schematic overview of the generation of HCT116 cell pools stably overexpressing BRIP1 WT or BRIP1 R162Q. BRIP1 WT or R162Q constructs contained C-terminal c-Myc and HA epitope tags. **b,** Representative Sanger sequencing traces confirming the presence of BRIP1 WT or BRIP1 R162Q in transduced HCT116 cells. **c, d,** Immunoblot analysis confirming expression of BRIP1 and Myc-tagged proteins in HCT116 cells expressing BRIP1 WT or R162Q, detected using BRIP1 and Myc antibodies. Vinculin served as a loading control. **e, j,** Functional characterization of HCT116 cells expressing BRIP1 WT or R162Q by colony formation assays following **e,** mitomycin C (MMC) (0.5;1; 2; 3; 4 μM/ml), **g,** ionizing radiation (IR) (2; 3; 4; 5 Gy), and **j,** hydroxyurea (HU) treatment 50;100;150;200;250 μM/ml). Representative colony plates (upper panels) and quantification of survival fractions (lower panels) are shown. Quantification was performed using EagleEye analysis software. Data represent the mean ± SD of three independent biological replicates.

### BRIP1^R162Q^ is mislocalized in response to DNA replication stress

Next, we studied the subcellular localization of the stably expressed HA-Myc-BRIP1^R162Q^ variant and the HA-Myc-BRIP1 wild-type counterpart in response to DNA replication stress induction by HU treatment. Of note, the predicted NLS of BRIP1 is localized at amino acids 159-174 ^6^. To this end, we performed immunofluorescence stainings of HU-treated cells using HA antibodies to detect the expressed HA-Myc-BRIP1^R162Q^ variant and HA-Myc-BRIP1 wild-type protein along with the DNA damage marker γH2AX to identify cells undergoing replication stress. Intriguingly, confocal microcopy revealed that the BRIP1^R162Q^ variant showed an altered nuclear distribution when compared to the wild-type protein (**Fig. 3a**). While wild-type BRIP1 showed strong pan-nuclear distribution, the BRIP1^R162Q^ variant signals were found to be reduced in cell nuclei in response to HU treatment. Quantification of the mean fluorescence intensity of HA-tagged wild-type and variant BRIP1 in γH2AX-positive nuclei revealed a profound and significant reduction in the intensity of the BRIP1^R162Q^ variant (**Fig. 3b**). These results suggest a defective subcellular distribution of the BRIP1^R162Q^ variant protein when compared to its wild-type counterpart. Moreover, whereas wild-type BRIP1 demonstrated broad colocalization with γH2AX in HU-treated cells, the BRIP1^R162Q^ variant only weakly colocalized with γH2AX (**Fig. 3c**), supporting our conclusion that the BRIP1^R162Q^ variant is mislocalized, or fails to be recruited, upon replication stress.

**Figure 3.**
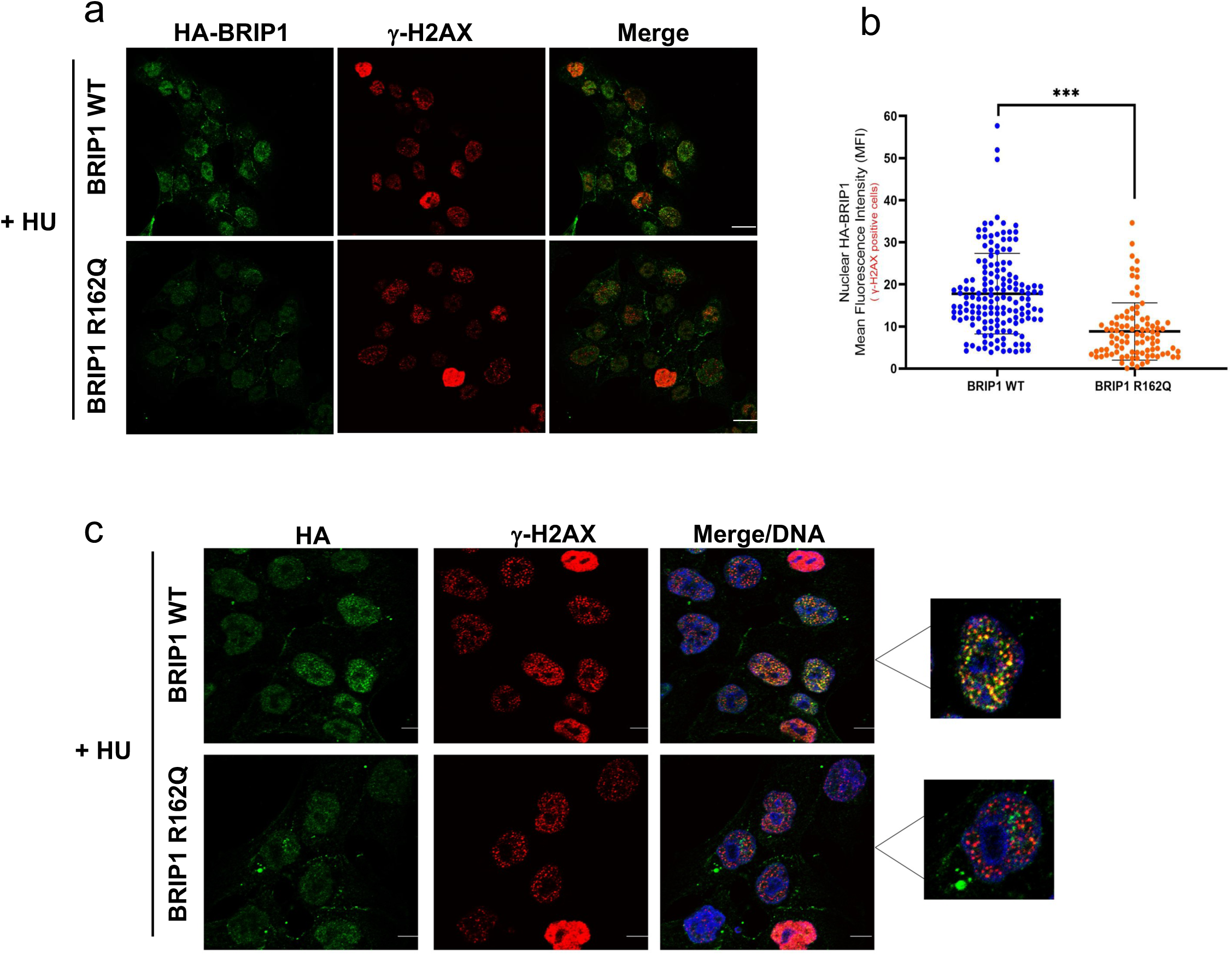
The BRIP1^R162Q^ variant shows an altered subcellular localization upon DNA replication stress. **a, c,** Immunofluorescence staining for HA-BRIP1 and HA-BRIP1 R162Q (anti-HA) and γH2AX in stably expressing cells upon overnight HU treatment (10 mM). Representative confocal microscopy images are shown (scale bar, 10 µm). **b,** Quantification of nuclear BRIP1, shown as mean fluorescence intensity (MFI) of HA-BRIP1 signal per nucleus in γH2AX-positive cells (defined as nuclei containing ≥5 γH2AX foci), measured with Image J. BRIP1 R162Q expressing cells show a 50% reduction in the MFI of BRIP1 in γH2AX-positive cells compared to BRIP1 WT expressing cells. A minimum of 100 cells per condition were analyzed across 2 independent experiments. Data are presented as mean ± SEM. Statistical analysis was performed using paired two-tailed t-tests (**p < 0.01 for BRIP1 foci recruitment; ***p < 0.001 for colocalization).

### BRIP1^R162Q^ variant expression triggers chronic DNA replication stress

Since BRIP1^R162Q^ variant cells are hyper-sensitive to DNA replication stress and show an altered subcellular distribution, we hypothesized that BRIP1^R162Q^ variant expressing cells may suffer from chronic DNA replication stress (in the absence of treatments). In line with our hypothesis, untreated BRIP1^R162Q^ cells showed increased phosphorylated replication protein A (pRPA) foci formation when compared to wild-type cells (**Fig. 4a,b**) indicating more ssDNA stretches occupied by RPA, which is a marker for replicating cells. To investigate activation of a chronic DNA replication stress response we performed DNA fiber assays were we pulse-labeled the cells with chlorodeoxyuridine (CldU) and iododeoxyuridine (IdU) and performed fiber spreading (**Fig. 4c**). Intriguingly, DNA fiber analysis indicated activation of chronic DNA replication stress in the untreated BRIP1^R162Q^ expressing cells characterized by less ongoing forks and an increase in terminated and stalled replication forks when compared to the wild-type counterparts (**Fig. 4d**). In addition, we also determined if BRIP1^R162Q^ expression affects DNA replication speed. BRIP1^R162Q^ expression resulted in a significantly reduced DNA replication speed (**Fig.4e**). Finally, we determined sister fork symmetry, which is a measure for the proper, synchronously coordinated propagation of the DNA replication fork^40^. Strikingly, BRIP1^R162Q^ expression resulted in an elevated sister fork asymmetry (**Fig. 4f**). Taken together, our results identify that BRIP1^R162Q^ expression leads to chronic DNA replication stress, which is a major driving force of genomic instability.

**Figure 4.**
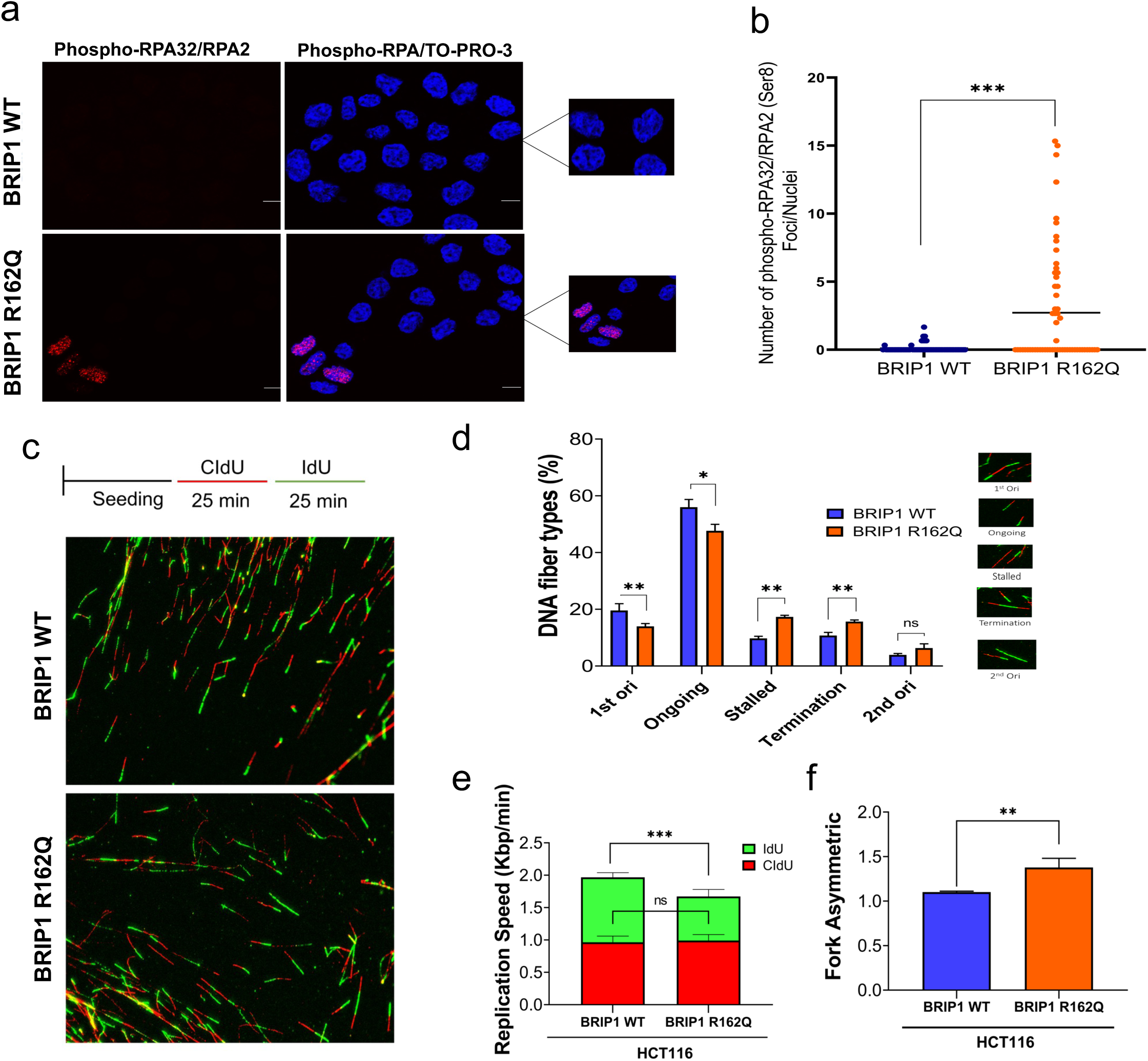
The BRIP1 R162Q variant triggers chronic DNA replication stress. Increased basal level of phospho-RPA32/RPA2 in cells expressing BRIP1^R162Q^. **a,** Cells were subjected to immunofluorescence staining with phospho-Ser8-RPA32/RPA2 antibody. Representative confocal microscopy images are shown (scale bar, 10 µm). Cells containing at least one distinct pRPA focus were scored as positive. **b,** Scatter plot showing phospho-RPA foci quantification. Quantification represents the number of pRPA-positive cells per nucleus (mean ± SEM) from *n* = 3 independent experiments. Statistical significance was assessed using a two-tailed paired *t* test. **c,** DNA fiber assay revealed altered replication dynamics in cells expressing the BRIP1 R162Q patient specific variant. Representative DNA fiber images from HCT116 cells stably expressing BRIP1 WT or BRIP1 R162Q, sequentially labeled with CldU (red) and IdU (green). **d,** Quantification of DNA fiber types. Representative schematics of each DNA fiber type are shown in the legend to the right of the graph. **e,** Replication fork progression was assessed. Cells were sequentially incubated with CldU and IdU for 25 min each. DNA fibers were spread, fixed, and immunostained with rat anti-BrdU antibody (detecting CldU) and mouse anti-BrdU antibody (detecting IdU), followed by appropriate secondary antibodies. Dual-labeled tracks were imaged by confocal microscopy, and red (CldU) and green (IdU) tracts were measured directly on the original confocal images using the overlay function of the LSM Image Browser software. Track lengths were determined in micrometers (µm) and converted to kilobases per minute (kb/min) using a factor of 2.59 kb per 1 µm. **f,** Fork asymmetry analysis, calculated as the ratio of the longer to the shorter IdU tract in bidirectional (single origin) replication fibers. All panels were derived from *n* = 3 independent biological replicates. Data represent mean ± SEM. Statistical analysis was performed using a paired two-tailed Student’s *t*-test; significance is indicated as \**p < 0.001, p < 0.01, p < 0.05, ns = not significant*.

### BRIP1^R162Q^ expression triggers genomic instability

To find out whether BRIP1^R162Q^ expression increases genomic instability, we first analyzed whether these cells show elevated levels of γH2AX foci, a marker of DSBs. Indeed, γH2AX foci numbers were increased in BRIP1^R162Q^ expressing cells (**Fig. 5a,b**), suggesting increased presence of DNA breaks, which may drive genomic instability. To directly assess the role of BRIP1^R162Q^ genomic instability we performed metaphase spreads and analyzed chromosomal aberrations in cells expressing wild-type and the BRIP1 variant (**Fig. 5c**). Strikingly, BRIP1^R162Q^ expressing cells showed a marked increase in the number of chromosomal aberrations when compared to the wild-type counterparts (**Fig. 5d**), indicative for a direct role of the variant in driving genomic instability. Detailed analysis of chromosomal aberrations revealed a pronounced increase in double-chromatid breaks and ring-shaped chromosomes, whereas single-chromatid breaks numbers were less prominent in the variant expressing cells (**Fig. 5e**). These results indicate that expression of the BRIP1^R162Q^ variant increases chromosomal aberrations and provokes a shift from single-stranded to double-stranded chromatid breaks.

**Figure 5.**
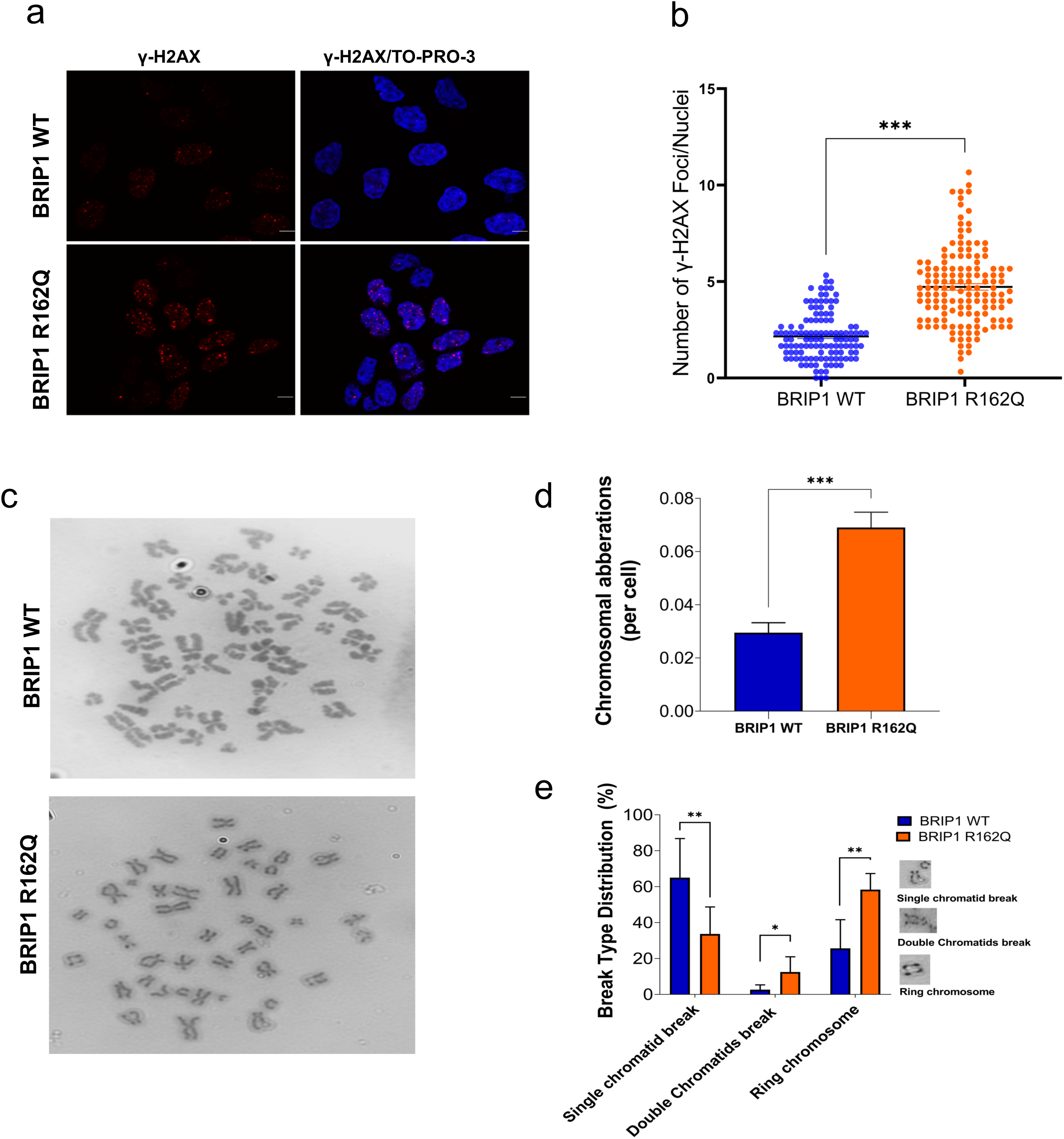
Pediatric cancer-linked BRIP1 R162Q variant stimulates genomic instability. **a,** BRIP1 R162Q expression increases γH2AX foci numbers. Immunofluorescence staining of phosphorylated histone H2AX (γH2AX) in untreated cells stably expressing BRIP1 WT or R162Q. Representative confocal microscopy images are shown (scale bar, 10 µm). **b,** Scatter plot showing γH2AX foci numbers in cell nuclei expressing BRIP1 R162Q compared to BRIP1 WT (**p < 0.01). At least 100 cells per condition were analyzed. Quantification is shown as mean ± SEM from n = 3 independent experiments. Statistical analysis was performed using paired two-tailed t-tests. **c,** BRIP1 R162Q expression stimulates genomic instability. Representative images of metaphase chromosome spreads from HCT116 cells stably expressing BRIP1 WT or BRIP1 R162Q. **d,** Quantification of chromosomal aberrations per cell. 30 metaphases have been investigated per replicate. BRIP1 R162Q cells showed a significant increase in chromosome aberrations compared to WT (paired two-tailed t-test). **e,** Distribution of chromosomal aberration types including single chromatid breaks, double chromatid breaks, and ring chromosomes shown as percentages of total aberrations. BRIP1 R162Q cells displayed an increased frequency of ring chromosomes compared to BRIP1 WT. Representative examples of each aberration type are shown to the right. Bars represent mean ± SEM from three independent experiments. *, p < 0.05; **, p < 0.01; ***, p < 0.001.

### BRIP1^R162Q^ expression results in increased G-quadruplex (G4) and R-loop formation, which is relieved by RNAseH1 expression

Previous studies identified an important role of BRIP1/FANCJ in unwinding G4 secondary DNA structures^11, 13^. To determine the impact of the cancer-linked BRIP1 variant on G4 load, we performed BG4-based immunostaining in the BRIP1^R162Q^ variant and BRIP1 wild-type cells. Our results revealed a strong increase in the G4 load in BRIP1^R162Q^ variant expressing cells (**Fig. 6a,b**).

**Figure 6.**
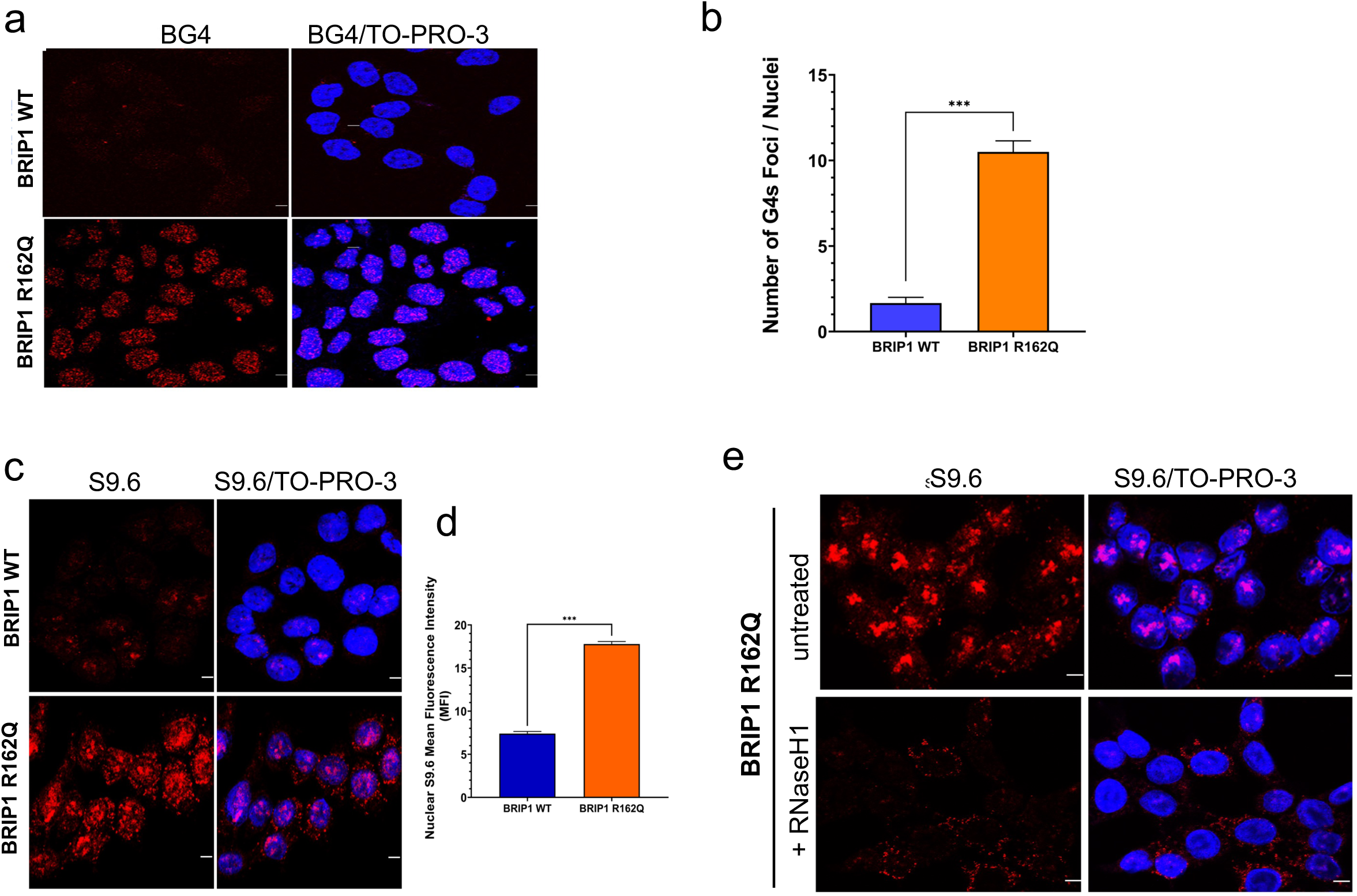
BRIP1 R162Q variant expressing cells show increased formation of G-quadruplex structures (G4s) and R-loops. **a,** Representative confocal microscopy images of HCT116 cells stably expressing BRIP1 WT or BRIP1 R162Q, proceed by indirect immunofluorescence staining with the G-quadruplex–specific BG4 antibody. BG4 was detected through sequential incubation with anti-FLAG and fluorophore-conjugated secondary antibodies. Nuclei were stained with TO-PRO-3. Scale bar, 20 µm. **b,** Quantification of nuclear G4 foci numbers in BRIP1 WT and BRIP1 R162Q expressing HCT116 cells. Minimum number of 100 nuclei analyzed per condition. Data represent the mean ± SEM from three independent experiments. two-tailed paired t-test was performed (**p < 0.001). **c,** Increased load of R-loop formation in BRIP1 R162Q variant expressing cells. Representative confocal microscopy images of HCT116 cells expressing BRIP1 WT or BRIP1 R162Q. **d,** Quantification of nuclear R-loop, shown as mean fluorescence intensity (MFI) of S9.6 signal per nucleus, measured with ImageJ. At least 150 nuclei per condition were analyzed in each replicate (n = 3 independent experiments). Data are presented as mean ± SEM. Statistical significance was determined by two-tailed paired t-test (**p < 0.001). **e,** Treatment with 10 U of recombinant RNaseH1 per coverslip abolished the S9.6 signal. R-loop staining was performed using the S9.6 antibody. Cells were fixed with cold methanol/acetone, and nuclei were counterstained with TO-PRO-3. Scale bar, 20 µm.

Interestingly, there is a close linkage between G4s and the formation of RNA:DNA hybrids termed R-loops has been demonstrated: G4 formation is regulated in a highly coordinated manner at Guanidine nucleotide-rich regions in the genome, and G4s are stabilized by formation of R-loops on the displaced ssDNA strand opposing the G4, termed G-loops^13^. Thus, we used S9.6 antibody-based immunofluorescence staining to analyze the R-loop levels in BRIP1^R162Q^ variant and wild-type cells. Intriguingly, we found that R-loop formation was strongly elevated in BRIP1^R162Q^ cells (**Fig. 6c,d**). RNAseH1 treatment, which resolves R-loops, confirmed the specificity of the S9.6 antibody staining (**Fig. 6e**).

To determine a potential causal relationship between increased formation of R-loops and G4s in BRIP1^R162Q^ expressing cells, we ectoptically expressed GFP-tagged RNaseH1 or GFP as a control and confirmed comparable transfection efficiencies using direct immunofluorescence analysis (**Fig. 7a**) and immunoblotting (**Fig. 7b**). Next, we analysed the effects of RNaseH1 expression on R-loop levels in BRIP1^R162Q^ expressing cells and, as expected, a strong reduction in R-loop load was evident (**Fig. 7c,d**). Next, we aimed to determine the effects of RNaseH1-mediated R-loop resolution on the G4 load in BRIP1^R162Q^ expressing cells. Intriguingly, analysis of G4 levels indicated a strong reduction of G4 levels upon RNaseH1 expression (**Fig. 7e, f**), suggesting a direct link between R-loops and G4s, potentially through G-loops^13^. In sum, our results indicate that BRIP1^R162Q^ expression provokes increased formation of G4s and R-loops, which can be reversed by RNaseH1 expression.

**Figure 7.**
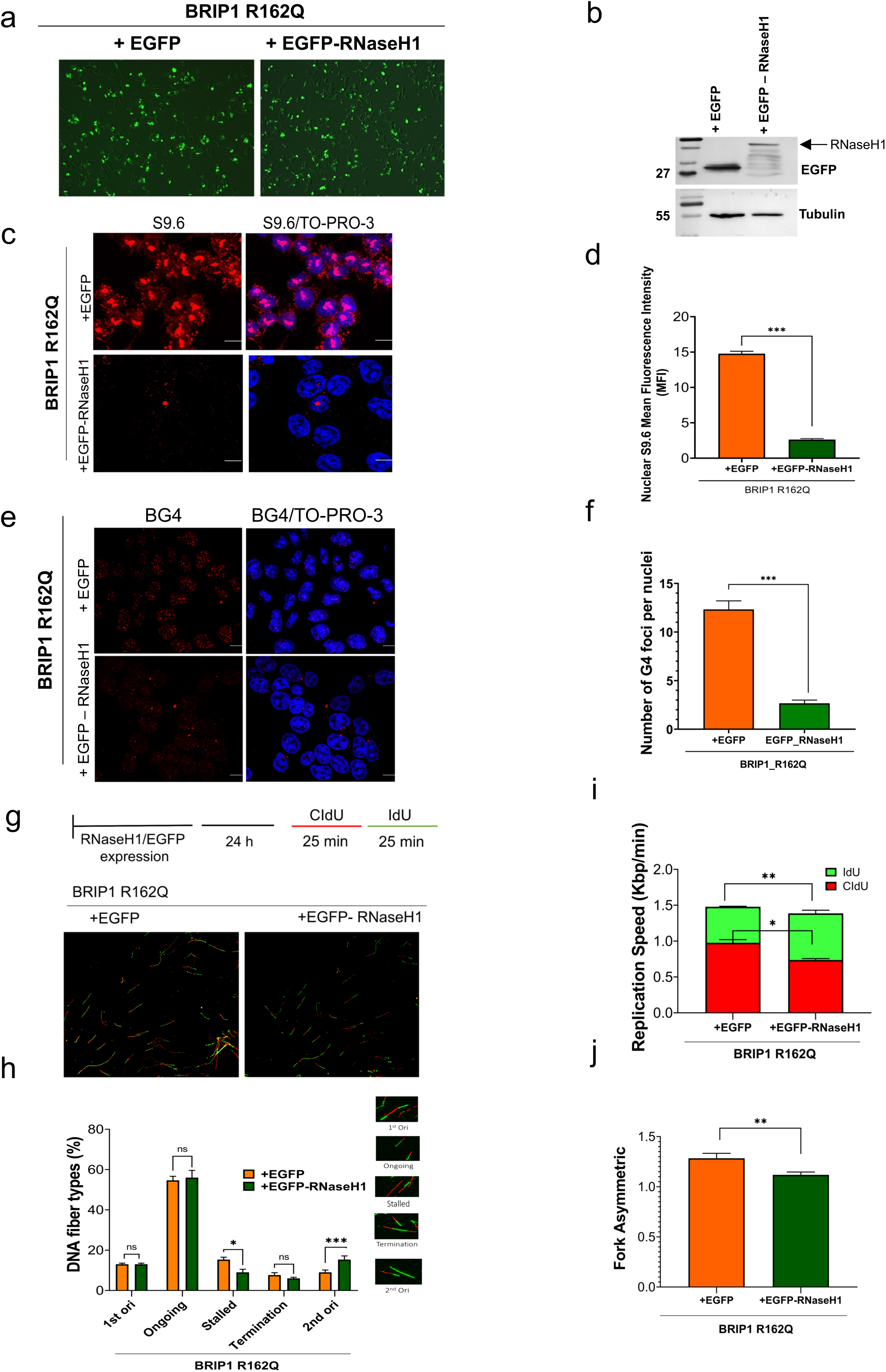
Resolving R-loops by RNaseH1 expression results in reduction of G4 structures in BRIP1 R162Q variant cells. **a,** HCT116 cells stably expressing BRIP1 R162Q were transiently transfected using PEI with empty vector (EV-EGFP) or RNaseH1-WT-eGFP. Transfection efficiency was assessed via EGFP fluorescence. **b,** immunoblot analysis confirming expression of GFP-tagged constructs in transfected cells. **c,** Immunofluorescence detection of R-loops using the S9.6 antibody in cells transfected with EGFP or EGFP-RNaseH1. **d,** Quantification of S9.6 mean fluorescence intensity (MFI) per nucleus. Data represent mean ± SEM of three independent experiments. **e,** Immunofluorescence detection of G-quadruplex (G4) structures using the BG4 antibody in cells transfected with EGFP or EGFP-RNaseH1. **f,** Quantification of BG4 fluorescence intensity per nucleus. Data represent mean ± SEM of three independent experiments. **g,** Representative DNA fiber assay images from HCT116 cells harboring BRIP1 R162Q transfected with empty vector (EV-eGFP) or RNaseH1-WT. DNA replication tracts were sequentially labeled with CldU (red) and IdU (green). **h,** Quantification of DNA fiber types. RNaseH1-WT expression reduced the proportion of stalled replication forks in cells harboring BRIP1 R162Q. **i,** While the overall fork speed was not significantly increased upon RNaseH1-WT expression, tracts displayed a more balanced distribution of CldU and IdU incorporation. **j,** Fork asymmetry analysis. RNaseH1-WT expression reduced fork asymmetry, indicating that left- and right-moving sister forks progressed at more similar speeds. Labeled tracks were visualized by confocal microscopy, and tract lengths were measured on raw LSM images. Lengths (µm) were converted to replication speed (kb/min) using a factor of 2.59 kb/µm. All data represent mean ± SEM from three independent experiments. At least 150 fibers per condition were analyzed in each replicate. Statistical significance was determined using paired two-tailed Student’s t-test (*p < 0.05, **p < 0.01, ***p < 0.001, ns = not significant). **c, e,** Representative confocal images are shown.

### Resolution of R-loops and G4s relieves DNA replication stress induced by BRIP1^R162Q^

To investigate potential causal relationship between R-loops and associated G4s and the chronic, steady-state replication stress response observed in BRIP1^R162Q^ expressing cells, we performed DNA fiber assays in BRIP1^R162Q^ cells expressing GFP-RNaseH1 or GFP as control (**Fig. 7g**). DNA fiber analysis revealed that RNaseH1 expression resulted in an increase in ongoing forks, accompanied by decreased fork stalling and increased 2nd origin firing (**Fig.7h**), indicative for reduced DNA replication stress. We did not observe an increase in DNA replication speed upon RNaseH1 expression in BRIP1^R162Q^ cells, which is presumably due to transfection-induced stress (**Fig. 7i**). However, strikingly, sister fork symmetry was significantly improved upon RNaseH1 expression in BRIP1^R162Q^ cells (**Fig.7j**), demonstrating harmonization of DNA replication fork progression. In sum, these results suggest a causal role of R-loops and presumably also of G4s and G-loops in triggering chronic replication stress in in BRIP1^R162Q^ expressing cells.

### BRIP1^R162Q^ expression generates vulnerabilities to pharmacological ATR and DNA-PK inhibition and G4-ligand Pyridostatin

DNA replication stress in cancer cells generates vulnerabilities which can be exploited by targeted therapeutic drugs ^41^. To this end, we pharmacologically inhibited the DNA damage checkpoint kinase ATR, which is activated in response to DNA replication stress and coordinates the replication stress response^42^. Strikingly, BRIP1^R162Q^ expressing cells were hypersensitive to the ATR inhibitor (ATRi) VE-822, when compared to the wild-type counterparts (**Fig. 8a,b**). BRIP1 KO cells showed an even higher sensitivity to ATRi (**Fig. 8a,b**), indicating that pertubation of BRIP1 function results in increased ATR dependency. In addition, BRIP1^R162Q^ expressing cells also showed an increased sensitivity to inhibition of DNA-PKcs (**Fig. 8c,d**), a kinase essential for repair of DNA double-strand breaks by the NHEJ pathway^43^. BRIP1 KO cells showed an even higher sensitivity to DNA-PK inhibition (**Fig. 8c,d**), indicating strong dependency of these cells on DNA-PK function.

**Figure 8.**
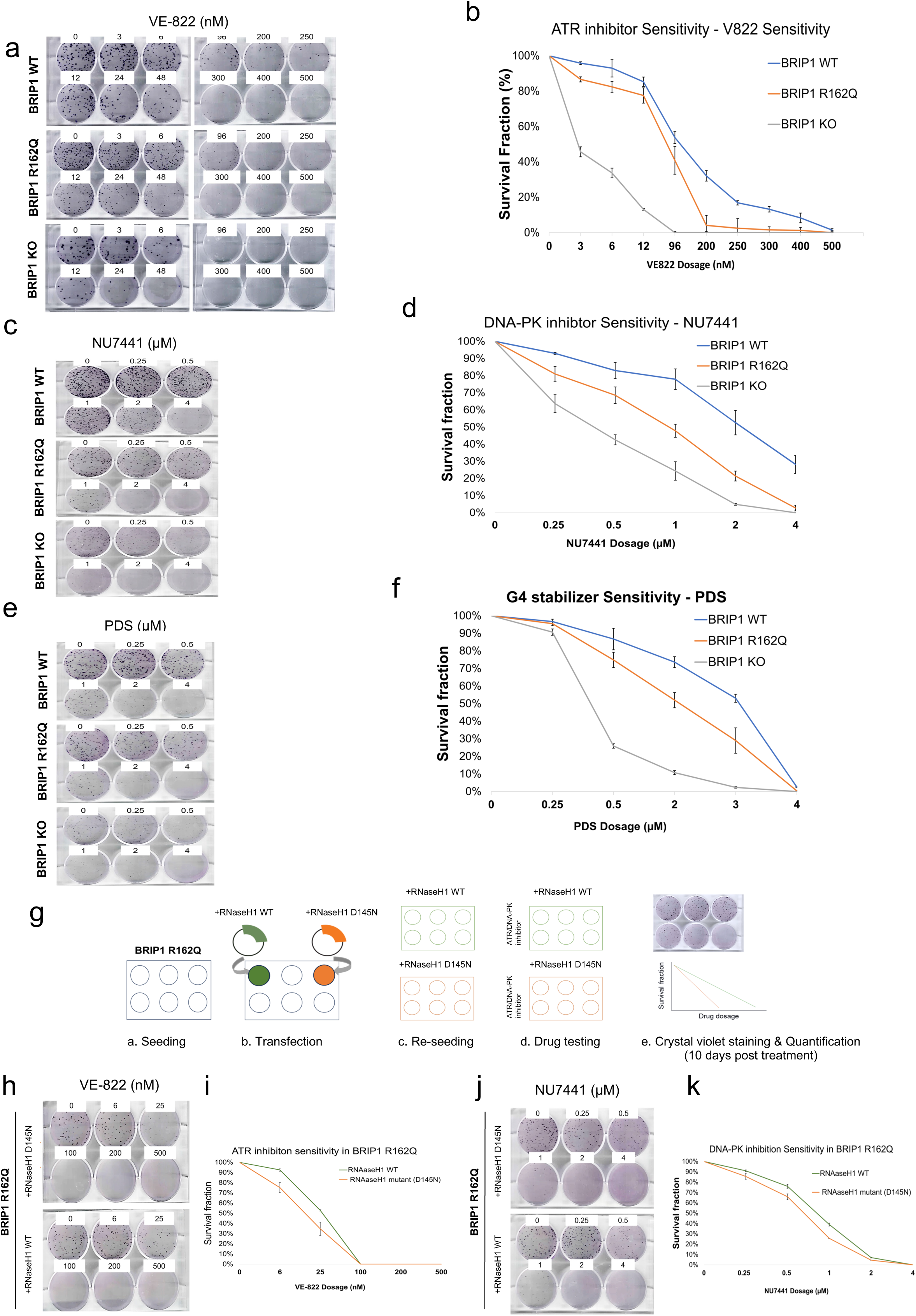
Increased sensitivity of BRIP1 R162Q expressing cells to DNA damage checkpoint kinase ATR inhibition, DNA-PK inhibition and G4-stabilizer Pyridostatin. **a,** BRIP1 R162Q cells show increased sensitivity to ATR inhibition. Representative colony formation assays and quantification of HCT116 cells expressing BRIP1 WT, BRIP1 R162Q, or BRIP1 knockout (KO) treated with increasing doses of the ATR inhibitor VE-822. Cells were treated overnight and colonies were allowed to form for 10 days prior to crystal violet staining. **b,** Quantification of survival fraction following ATR inhibition by VE-822. **c,** BRIP1 R162Q expressing cells show increased sensitivity to DNA-PK inhibition. Representative colony formation assays and quantification following overnight treatment with increasing doses of the DNA-PK inhibitor NU7441. **d,** Quantification of survival fraction following DNA-PK inhibition using NU7441 **e,** BRIP1 R162Q cells show increased sensitivity to treatment with G4-stabilizer Pyridostatin. Representative colony formation assays following overnight treatment with increasing concentrations of Pyridostatin (PDS) are shown. After drug removal, cells were cultured for 10 days before fixation and staining. **f,** Quantification of surviving fraction following Pyridostatin treatment **b,d,f**, Quantification was performed using EagleEye analysis software. Data represent the mean ± SD of three independent biological replicates. **g,** Schematic representation of the workflow used to perform colony formation assays in BRIP1R162Q cells upon R-loop removal by ectopic RNaseH1 expression. BRIP1 R162Q cells were transfected with RNase H1 WT or catalytically inactive RNaseH1 D145N, re-seeded, and treated the next day with the drugs indicated. Colonies were stained with crystal violet after 10 days**. h-k,** RNaseH1 expression decreases sensitivity to ATR VE-822 (**h,i,**) and DNA-PK NU7441 (**j,k,**) inhibition in BRIP1 R162Q expressing cells. **h,j,** Show representative colony formation plates and **i, k,** show quantifications of clonogenic survival. Data represent mean values ± SEM from three biologically independent experiments. Colony formation was quantified using EagleEye analysis software.

Since BRIP1 has an important role in unwinding G4 DNA secondary structures, we also tested sensitivity towards the G4 ligand Pyridostatin, which stabilizes G4s and has been shown to trigger cell death in cancer cells^44^. Strikingly, BRIP1^R162Q^ expressing cells exhibited increased sensitivity to Pyridostatin treatment (**Fig. 8e,f**). BRIP1 KO cells also demonstrated hypersensitivity to the G4 ligand (**Fig. 8e,f**). Taken together, our results indicate that BRIP1^R162Q^ expression provides specific vulnerabilities which can be exploited by targeted cancer therapeutics.

### Resolving R-loops by RNAseH1 decreases sensitivity of BRIP1^R162Q^ expressing cells to ATR and DNA-PK inhibition

To investigate whether the increased drug sensitivity to ATRi and DNA-PKi relies on the increased R-loop formation and G4 load in BRIP1^R162Q^ expressing cells, we transfected BRIP1^R162Q^ cells with GFP-RNaseH1, to dissolve R-loops, or catalytically-inactive GFP-RNaseH1^D145N^ as control, and re-seeded the cells to performed colony formation analysis (**Fig. 8g**). Intriguingly, colony formation analysis revealed that RNaseH1 expression, which resolves R-loops and G-loops, resulted in reduced ATRi sensitivity when compared to cells expressing catalytically-inactive GFP-RNaseH1^D145N^ (**Fig. 8h,i**). In addition, comparable results were obtained for cells treated with the DNA-PKi, where RNaseH1 expression, in contrast to catalytically-inactive GFP-RNaseH1^D145N^, resulted in reduced sensitivity against DNA-PKi (**Fig.8j,k**). Taken together, our results suggest that R-loops and presumably also G-loops, which are both resolved by RNaseH1^13^, contribute to therapeutic drug sensitivity of BRIP1^R162Q^ variant expressing cells.

## Discussion

Development of pediatric cancer is largely determined by inherited germline variants in cancer predisposition genes, with increasing evidence for a role of early-onset and adult-onset DNA repair-regulating genes^5, 34^. Efficient DNA repair plays a fundamental role in tumor suppression and the prevention of genomic instability, which is a major cause of cancer development. Accordingly, genes defining high-fidelity DNA double-strand break repair by homologous recombination (HR), including BRCA1, BRCA2 and Palb2, as well as the Fanconi Anemia (FA) pathway genes implicated in ICL have been frequently found to be altered in the germline of early-onset cancer syndromes^45^.

To evaluate the pathogenic potential of the identified BRIP1^R162Q^ variant, we characterized its impact on genome stability and cancer cell vulnerability.. BRIP1 is a 5’-3’-directed DNA helicase^15^ which functions in ICL, HR and stabilization of DNA replication forks during replication stress. Unexpectedly, our *in vitro* helicase assays identified BRIP1^R162Q^ as a hyper-active DNA helicase, indicating an altered enzymatic function of this variant. Of note, previous studies identified BRIP1 germline SNVs with reduced DNA helicase functions in women with early-onset breast cancer and in patients with ovarian cancer^15, 17^. To our knowledge our findings provide the very first link between a hyperactive, hypermorphic BRIP1 variant and cancer development. To study its role in a cellular model system, we generated HCT116 cell pools stably expressing HA-BRIP1^R162Q^ and the wild-type counterpart HA-BRIP1 wt. This model mimicks our pediatric cancer patient setting since the BRIP1 variant was monoallelic and expressed along with a BRIP1 wild-type allele. Our sensitivity tests indicated specific sensitivity of BRIP1^R162Q^ expressing cells to the DNA replication stress-inducing drug HU, while no altered sensitivity in response to IR or treatment with the DNA crosslinker MMC was detectable. Interestingly, we found that BRIP1^R162Q^, when compared to its wild-type counterpart, is mislocalized in response to replication stress -or impaired in proper recruitment-suggesting that BRIP1^R162Q^ may have comprimized activity in unwinding its *bona fide* substrates, such as G4 DNA secondary structures. In line with this hypothesis, our findings revealed that BRIP1^R162Q^ expression resulted in chronic DNA replication stress, and in an increase in chromosomal aberrations, indicating a link between BRIP1^R162Q^-induced replication stress and increased genomic instability, which is major driving force for cancer development. These findings suggest a causal relationship between the BRIP1^R162Q^ germline variant and genome instability, arguing for a direct role in pediatric and early-onset cancer development.

In addition, our findings indicate that BRIP1^R162Q^ expression evokes a strong increase in G4 structures and R-loops in the cell nucleus. Both G4’s and R-loops can be recognized as DNA lesions^29^ and are established activators of the DNA replication stress and DNA damage checkpoint kinase ATR, and also of ATM due to replication-evolved DSBs^46^. Remarkably, ectopic expression of RNaseH1, which resolves R-loops^29^, also resulted in a strong reduction of G4 structures in the BRIP1^R162Q^ expressing cells, and relieved DNA replication stress in the cells. These results suggest a causal relationship between increased R-loop and G4 formation in provoking DNA replication stress in in the BRIP1^R162Q^ expressing cells. Furthermore, recent evidence indicates a direct link between G4’s and R-loops, since R-loops form on the displaced DNA strand opposing the G4, termed G-loops^13^, through a highly-coordinated G-loop assembly involving the G4, RPA, ATR, BRCA2 and Rad51 along with a complementary RNA bound to hnRNPA1^13^. Based on this study, we postulate that reduction of G4 levels in BRIP1^R162Q^ expressing cells through RNaseH1 expression likely stems from resolution of R-loop moieties of G-loops, which has been shown to destabilize G4 structures^13^. Intriguingly, BRIP1/FANCJ was shown to shape the G4 genomic landscape, and FANCJ deficiency results in elevated G4 levels and increased genome instability^13^, indicating that balanced G-loop assembly and disassembly is fundamental to maintain genomic stability. Thus, the deregulated BRIP1/FANCJ function in BRIP1^R162Q^ expressing cells likely alters the G4 genomic landscape, and thereby increases genomic instability through induction of chronic DNA replication stress. Future studies are required to address this highly interesting link. In addition, it will be important in the future to unravel the underlying mechanisms by which helicase hyperactivity and/or mislocalisation contribute to the genomic instability phenotype upon BRIP1^R162Q^ expression.

Our findings suggest the following mechanistic framework by which the childhood cancer-linked BRIP1^R162Q^ germline variant participates in cancer development (**Fig. 9a**): BRIP1^R162Q^ encodes a hyper-active and mislocalized DNA helicase enzyme, which stimulates increased formation of G4’s, R-loops and presumably G-loops by unbalancing BRIP1/FANCJ function in G-loop disassembly. Increased R-loop and G4 accumulation stimulates chronic DNA replication stress by stalled replication forks and fork assymmetry, which gives rise to increased DSBs and genomic instability (**Fig. 9a**), which is a major driving force for cancer development.

**Figure 9.**
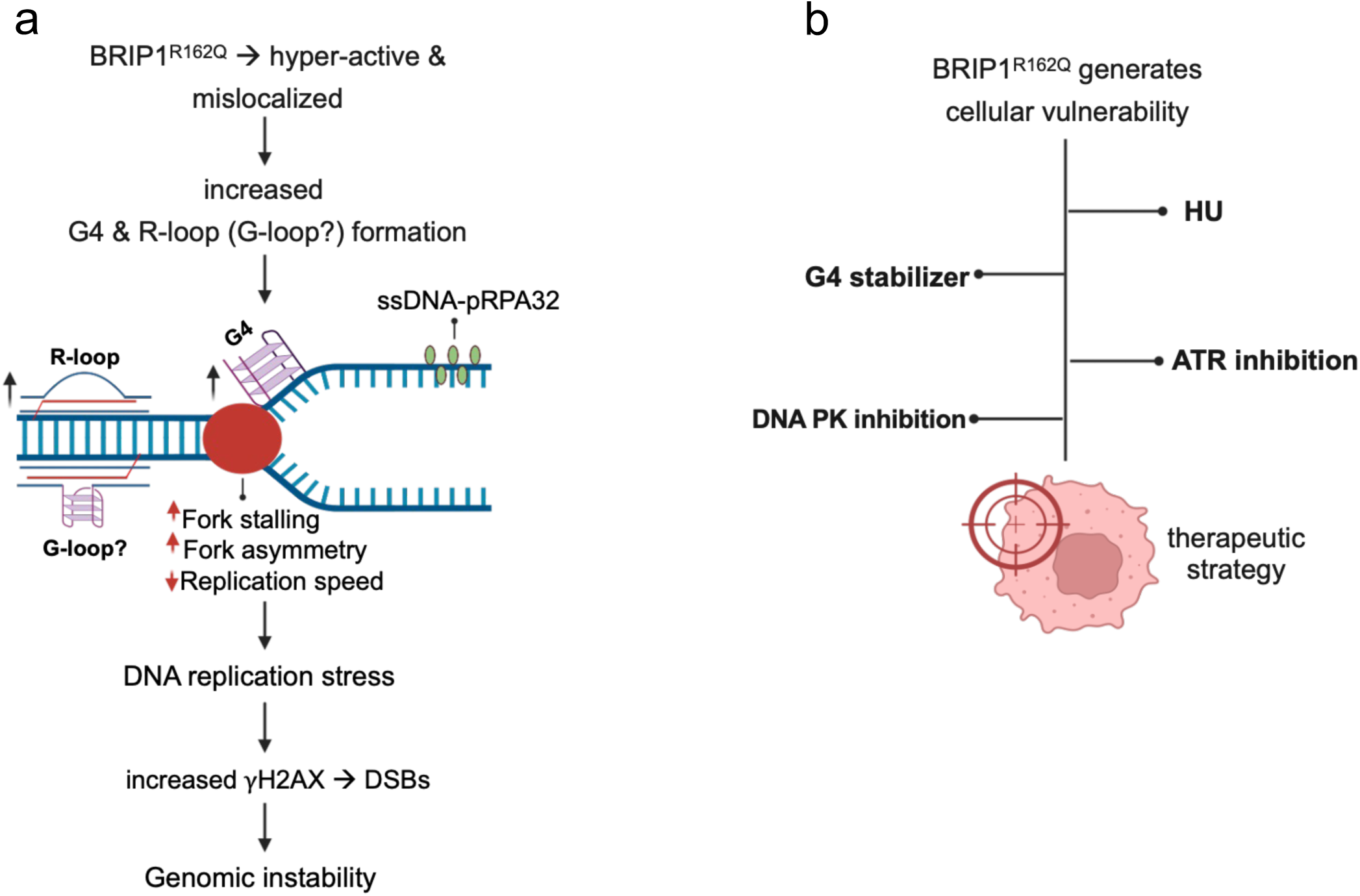
Proposed mechanistic framework for the role of BRIP1^R162Q^ in childhood cancer development and therapeutic strategies for targeting vulnerabilities. **a,** Proposed mechanistic framework for the role of BRIP1^R162Q^ variant in childhood cancer development. For details, see discussion part. **b,** Potential therapeutic strategies to exploit vulnerabilities of BRIP1^R162Q^ variant expressing cancer cells. For details, see discussion part. Graphical illustrations were generated using BioRender.

Our work also implies potential therapeutic strategies to exploit specific vulnerabilities identified in BRIP1^R162Q^ expressing cancer cells (**Fig. 9b**). Our results identified induction of chronic DNA replication stress and an increased G4 load are tighly linked to BRIP1^R162Q^ variant expression. Along these lines, our results indicate that BRIP1^R162Q^ expression sensitizes cancer cells to therapeutic induction of DNA replication stress by HU treatment. In addition, cells expressing BRIP1^R162Q^ show increased sensitivity to pharmacological inhibition of the DNA damage checkpoint kinase ATR, which is activated in response to DNA replication stress^42^. Moreover, hypersensitivity of BRIP1^R162Q^ expressing cancer cells to inhibition of DNA-PK, an essential regulator of DSB repair by the NHEJ pathway^43^, was also identified. Finally, BRIP1^R162Q^ expressing showed hypersensitivity to treatment with the G4-stabilizing ligand Pyridostatin, a drug previously shown to show high efficacy on the killing of cancer cells characterized by elevated G4 levels^44, 47^. Taken together, our results identify a mechanistic framework suggesting a function of the pediatric cancer-linked BRIP1^R162Q^ variant in genomic instability and cancer development, and identify specific vulnerabilities of variant expressing cancer cells which can be exploited by targeted cancer therapy.

## Methods

### Whole Exome Sequencing (WES) analysis

Genetic analyses were performed within the prospective study “*Germline Mutations in Children with Cancer*”^48, 49^. Children with newly diagnosed malignancies and their parents were offered trio-based whole-exome sequencing (WES) independent of family history or clinical suspicion of a cancer predisposition syndrome. Clinical, demographic, and family history data, including three-generation pedigrees, were systematically collected. Written informed consent was obtained from all participants or their legal guardians following at least two dedicated genetic counseling sessions, and families were given sufficient time for consideration prior to enrollment. The study was approved by the Ethics Committee of Heinrich Heine University Düsseldorf (vote no. 4886R) and conducted in accordance with the Declaration of Helsinki. Genomic DNA was extracted from peripheral blood of the index patient and both parents. Exome capture was performed using the SureSelect Human All Exon V5+UTR kit (Agilent Technologies) and paired-end sequencing was conducted on Illumina platforms to achieve a mean on-target coverage of ≥80×. Reads were aligned to the GRCh37 reference genome using BWA-MEM, followed by duplicate marking and variant calling with GATK and VarScan2. Variant annotation included Ensembl Variant Effect Predictor (VEP) and functional prediction by SIFT, PolyPhen-2, and CADD. Filtering focused on rare (minor allele frequency <1%) protein-altering variants, with additional restriction to genes implicated in cancer predisposition and DNA damage response pathways. Variants were prioritized based on rarity, predicted deleteriousness, inheritance pattern, and biological plausibility. In the index patient with metastatic osteosarcoma, a maternally inherited *BRIP1* missense variant, NM_032043.3:c.485G>A, located within the nuclear localization signal, and a paternally inherited *HIPK2* missense variant, p.(Thr602Pro), were identified. The *BRIP1* variant was initially considered deleterious based on *in silico* predictions, and both variants were discussed in the context of a putative digenic effect impacting DNA damage response signaling^30^. The patient’s mother, who carried the *BRIP1* variant, was diagnosed with breast cancer at 46 years of age. According to current ACMG/AMP-based interpretation frameworks integrating allele frequency, gene–disease validity, computational evidence, segregation data, and phenotype relevance, the *BRIP1* variant remains classified as a variant of uncertain significance. It is listed in ClinVar as of uncertain significance (SCV000184600, SCV000689394; accession VCV000141318.30) and occurs at low frequency in population databases (gnomAD). ClinGen curation confirms *BRIP1* as a gene associated with autosomal-dominant ovarian cancer predisposition and recessive Fanconi anemia but not with hereditary breast cancer. Although *HIPK2* s functionally implicated in DNA damage response signaling, current publica curation resources do not support its classification as a hereditary cancer predisposition gene. The *HIPK2* variant is similarly rare in gnomAD an, in the absence of supporting clinical or functional evidence, is likewise best regarded as a variant of uncertain clinical significance.

### Preparation of expression vectors

The codon-optimized expression construct for purification of recombinant human FANCJ/BRIP1 (pFastBac1-FANCJ-STREP-WT) in insect cells (Sf9; *Spodoptera frugiperda*) was a kind gift by Julian Stingle^50^. This construct contained TwinStep tag at C-terminal of FANCJ preceded by TEV protease site. To prepare the variant pFB1-FANCJ-R162Q-STREP, the WT construct was mutagenized with the primers FANCJ_R162Q_F (GAAGCGTATCCAGCCTCTGGAAAC) and FANCJ_R162Q_R (TTCTCGACCTGGAAG) using the Q5 site-directed mutagenesis kit according to the manufacturer’s instructions. The correct sequence of pFB-FANCJ-STREP-WT and pFB-FANCJ-R162Q-STREP constructs were verified by full length plasmid sequencing by Eurofins.

### Recombinant protein expression and purification

To express FANCJ WT and R162Q mutant in insect cells, bacmid, primary and secondary baculoviruses were prepared according to the manufacturer’s instructions (Bac-to-bac system, Life technologies). To express recombinant FANCJ WT, 1.6 L *Sf9* insect cells were seeded at 0.5 million cells/ml and infected next day at 1 million/ml with FANCJ WT secondary baculovirus. The infected cells were incubated at 28 °C for 72 h with continuous agitation. Cells were collected by centrifugation at 500*g* for 10 min and washed once with 1× PBS (137 mM NaCl, 2.7 mM KCl, 10 mM Na2HPO4, 1.8 mM KH2PO4). The collected pellets were snap-frozen in liquid nitrogen and stored at −80 °C until further use. All subsequent steps were performed either on ice or at 4 °C. The cells pellets were resuspended in 3 volumes of lysis buffer containing 50 mM Tris-HCl pH 7.5, 1 mM ethylenediaminetetraacetic acid (EDTA), protease inhibitor cocktail (1:400), 30 µg /ml leupeptin (Merck), 1 mM phenylmethylsulfonyl fluoride (PMSF), 1 mM Tris(2 carboxyethyl) phosphine (TCEP) and incubated for 20 min with continuous agitation. Next, 50% glycerol and 3 M KCl were added sequentially to the final concentrations of 16.5% and 305 mM, respectively, and the suspension was further incubated for 30 min with continuous agitation. The suspension was centrifuged at ∼18,500 rpm for 30 min to obtain the soluble extract. The Strep Tactin 4Flow resin (IBA Lifesciences) was pre-equilibrated with strep wash buffer I (50 mM Tris-HCl pH 7.5, 500 mM KCl, 1 mM TCEP, 1 mM PMSF and 10% glycerol) and added to 50 ml tubes containing soluble extract. These tubes were subsequently incubated for 1 h with continuous rotation. After incubation, the resin was washed batchwise two times by centrifugation at 2,000*g* for two min and four times on column with strep wash buffer. Protein was eluted from the resin using strep elution buffer (50 mM Tris-HCl pH 7.5, 125 mM KCl, 1 mM TCEP, 1 mM PMSF and 10% glycerol supplemented with 2.5 mM desthiobiotin (IBA Lifesciences) and total protein was estimated using the Bradford assay. To remove the TwinStrep tag, 1/10 (w/w) of TEV protease to the total protein was added to strep eluate and incubated for 2 h with gentle agitation. Following cleavage of TwinStrep tag, the protein sample was loaded on a 1 mL HiTrap Heparin HP affinity column equilibrated in buffer A (50 mM Tris-HCl pH 7.5, 125 mM KCl, 1 mM TCEP, 1 mM PMSF and 10% glycerol) using AKTA pure (Cytiva Life Sciences) and eluted with salt gradient using buffer A and buffer B (50 mM Tris-HCl pH 7.5, 1 M KCl, 1 mM TCEP, 1 mM PMSF and 10% glycerol). Eluted fractions corresponding to the UV and conductivity peaks from AKTA were pooled, dialyzed overnight using the Pur-A-Lyzer Midi Dialysis Kit (Merck). Following overnight dialysis, fractions were concentrated using 100 kDa ultra centrifuge filter (Millipore) and snap-frozen in liquid nitrogen and stored at −80℃. The same purification procedure was used to purify FANCJ R162Q using 800 mL *sf9* insect cell culture.

### Preparation of DNA substrates and oligonucleotides used for in vitro analysis

All DNA oligonucleotides used in the in vitro analysis were commercially synthesized and purchased from Integrated DNA Technologies (IDT). To prepare various substrates used in this study, when needed, combination(s) of DNA oligonucleotides were annealed together by mixing and heating them at 95 °C for 3 min, followed by gradual cooling of the samples overnight unless indicated otherwise. To prepare 5’ overhang substrate for DNA unwinding assays, 5’-FAM labelled Oligo (Oligo 25nts) was annealed with oligo 2 in 1:1,2 ration respectively as described above. The G-quadruplex DNA substrates were formed by heating 10 µM of 5’FAM-Oligo 95_TPG4 in 100 mM Tris HCl, pH 7.5, 1mM EDTA, and 1M NaCl to 95°C for 5 min followed by incubation overnight at 60°C (20 hrs). The prepared G4 substrate was cooled down to 30°C and resolved by a 6% native polyacrylamide gel (19:1 acrylamide-bisacrylamide, Bio-Rad) using the Mini-Protean Tetra Cell electrophoresis system (Bio-Rad) at 100 V for 1 h. The FAM in oligos indicate the position of 6-carboxyfluorescein.

### DNA unwinding assay

The unwinding assays were performed in 15 µl helicase buffer containing 25 mM Tris-HCl pH 7.5, 2 mM ATP, 2 mM MgCl2, 1 mM DTT, 50 mM NaCl, 0.1 mg/ml bovine serum albumin (BSA, New England Biolabs), 1 mM PEP (phosphoenolpyruvate, Sigma-Aldrich), 10 U/ ml pyruvate kinase (Sigma-Aldrich) and 25 nM DNA substrate (in molecules). All steps except for the assembling reactions and protein addition were performed in the dark. The reactions were assembled on ice and recombinant proteins were added, mixed and incubated at 37 °C for 30 min. The reactions were stopped with 5 µl of 2% stop solution (0.2% SDS, 30% glycerol, 150 mM EDTA, bromophenol blue) and 1 µl proteinase K (Roche, 18.4 mg ml−1) by incubating the tubes for 10 min at 37 °C. The products were resolved by 10% native polyacrylamide gel (19:1 acrylamide-bisacrylamide, Bio-Rad) using the Mini-Protean Tetra Cell electrophoresis system (Bio-Rad) at 100 V for 1 h. The gels were directly imaged in ChemiDoc MP imaging system.

### G4-quadruplex unwinding assay

The G-quadruplex unwinding assays were performed in 15 µl buffer containing 25 mM Tris-HCl pH 7.5, 5 mM ATP, 4 mM MgCl2, 1 mM DTT, 50 mM NaCl, 0.1 mg/ ml bovine BSA, and 25 nM G-quadruplex DNA substrate (FAM-Oligo 95_TP-G4 oligo)^35^. These assays were carried out and processed as described for DNA unwinding assay above except the final products were resolved with 6% native polyacrylamide gel.

### DNA oligo used for biochemical assays

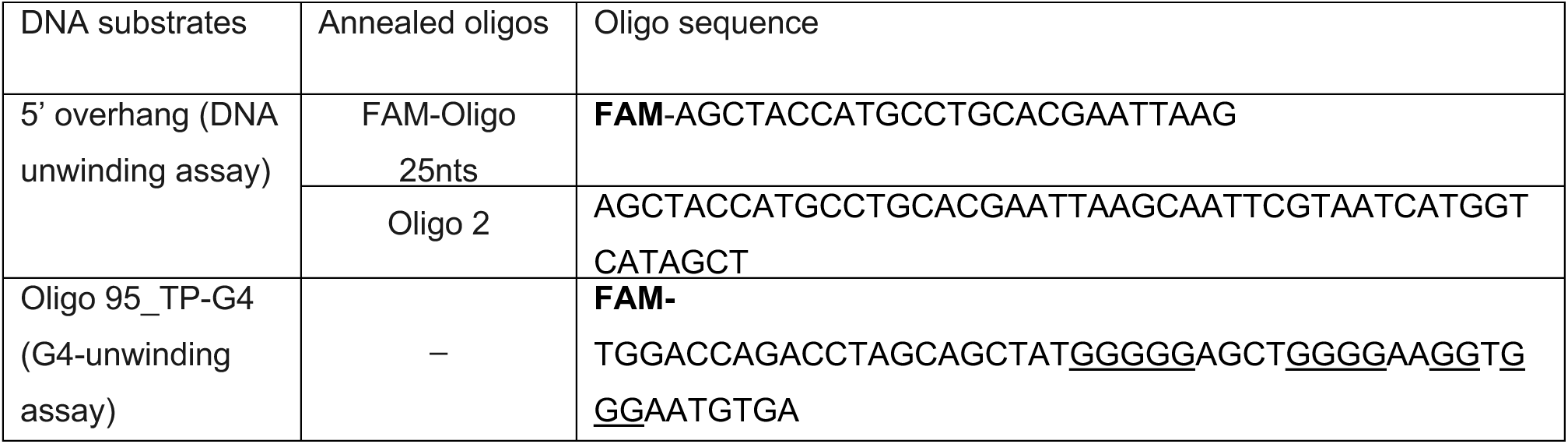

The **FAM** in oligos indicate the position of 6-carboxyfluorescein. The G-quadruplex forming guanines are underlined in the Oligo 95_TP-G4 oligo^35^.

### Cell culture and transfection

HCT116, HEK293T, and U2OS cells (all obtained from ATCC) were maintained in DMEM supplemented with 10% (v/v) fetal calf serum (FCS), 100 U/mL penicillin, 100 μg/mL streptomycin, 10 mM HEPES, and 2 mM L-glutamine at 37 °C in a humidified atmosphere with 5% CO₂. Transient transfections were performed using Lipofectamine 3000 (Thermo Fisher Scientific) according to the manufacturer’s instructions or by polyethylenimine (PEI)-mediated transfection. For PEI-based transfection, cells were seeded to ∼70–80% confluency and transfected with plasmid DNA using linear PEI (1 mg/mL stock) at a DNA:PEI ratio of 1:3. DNA–PEI complexes were formed in Opti-MEM by incubation for 30 min at room temperature and added to cells, followed by medium replacement after 6–8 h. Transfection efficiency was typically assessed 24–48 h post-transfection. For R-loop removal, cells were transiently transfected with an EGFP empty vector control (pEGFP-C1, Takara Bio, #632470), EGFP-tagged RNaseH1 wild-type (WT), or the catalytically inactive RNaseH1 D145N mutant as indicated. The RNaseH1 constructs (pEGFP-RNaseH1 WT, Addgene #108699; and RNaseH1 D145N-EGFP) were kindly provided by Brian Luke (Institute of Molecular Biology, Mainz). Cells were processed 24 h post-transfection for downstream assays as indicated.

### Chemotherapeutic drugs treatments

Cells were incubated with culture medium supplemented with mitomycin C (MMC; Carl Roth, 4150.1), hydroxyurea (HU; Sigma-Aldrich, H8627), pyridostatin (PDS; MedChemExpress, HY-15176), NU7441 (KU-57788; Selleckchem, S2638), or VE-822 (berzosertib; Selleckchem, S7102) at the indicated concentrations and time points. VE-822 was dissolved in DMSO; MMC, HU, PDS, and NU7441 were dissolved in sterile water. Control treatments were performed with the corresponding solvents. For colony formation assays, cells were treated with MMC (0–4 µM, 1-h pulse), HU (0–250 µM, continuous exposure), PDS (0–4 µM, overnight), NU7441 (0–4 µM, overnight), or VE-822 (0–500 nM, overnight). After treatment, drug-containing medium was replaced with fresh growth medium and cells were cultured under standard conditions. For immunofluorescence and western blotting, cells were treated with 10 mM HU overnight prior to fixation or lysis. For ionizing radiation experiments, HCT116 cells were treated using a Gammacell 2000 irradiator.

### Antibodies

Primary antibodies used for immunoblotting included α-tubulin (D3U1W, #86298T), HA (C29F4, #3724), ATM (D2E2, #2873), BRIP1/FANCJ (#4578), FLAG (DYKDDDDK, #14793), phospho-ATM (Ser1981, #4526), and phospho-Chk1 (Ser345, #2341S) from Cell Signaling Technology; Chk1 (#C9358) from Sigma-Aldrich; Myc (9E10, #sc-40) and Rad51 (3C10, #sc-53428) from Santa Cruz Biotechnology; phospho-ATR (Thr1989, #PA5-77873) from Life Technologies; phospho-Chk2 (Thr68, #2661T) from New England Biolabs; and phospho-histone H2A.X (Ser139, JBW301, #05-636) from Merck Millipore. The HIPK2 antibody (rb1) was obtained from Pineda (self-made). Secondary antibodies for immunoblotting were horseradish peroxidase–conjugated goat anti-mouse IgG (H+L, #115-005-146) and goat anti-rabbit IgG (H+L, #111-035-144) from Jackson ImmunoResearch. For immunofluorescence, primary antibodies included anti-G-quadruplex (BG4, #MABE917) and anti-DNA–RNA hybrid (S9.6, #MABE1095) from Sigma-Aldrich; FLAG (#14793) and HA (C29F4, #3724) from Cell Signaling Technology; phospho-histone H2A.X (Ser139, #05-636) from Merck Millipore; and phospho-RPA32 (Ser8, #54762) from Cell Signaling Technology. Secondary antibodies were Alexa Fluor 488–conjugated goat anti-mouse (#115-546-062, Jackson ImmunoResearch), Alexa Fluor 488–conjugated goat anti-rabbit (#A11070, Life Technologies), Cy3 goat anti-mouse (#115-166-006, Jackson ImmunoResearch), and Cy3 goat anti-rabbit (#111-166-003, Jackson ImmunoResearch). For DNA fiber assays, primary antibodies included mouse anti-BrdU (clone B44, #347580, BD Biosciences) and rat anti-BrdU (clone BU1/75, #ab6326, Abcam). Secondary antibodies were Cy3 donkey anti-rat (#712-165-150, Jackson ImmunoResearch) and Alexa Fluor 488 goat anti-mouse (#115-546-062, Jackson ImmunoResearch).

### Metaphase spread preparation and analysis of chromosomal aberrations

Cells were seeded in 6-well plates at ∼30% confluency 24 h before harvest. To enrich for mitotic cells, cultures were treated with colcemid (200 ng/mL) for 6 h at 37 °C. Cells were collected in complete medium, and pelleted by centrifugation. Cell pellets were resuspended in PBS and subjected to hypotonic treatment by dropwise addition of pre-warmed 0.075 M KCl to a final volume of 10 mL, followed by incubation for 10 min at 37 °C. Cells were then fixed by gradual addition of ice-cold methanol:acetic acid (3:1, v/v). After centrifugation, the fixation step was repeated twice with fresh fixative. Fixed cells were stored in fixative at −20 °C or processed directly for slide preparation. For metaphase spread preparation, fixed suspensions were dropped onto pre-warmed humidified glass slides and air-dried at room temperature. Metaphase spreads were visualized by microscopy using a 100× oil immersion objective. Chromosome aberrations were scored in metaphase cells under identical imaging settings across allexperimental conditions.

### Generation of BRIP1 knockout cell lines

Single guide RNAs (sgRNAs) targeting *BRIP1* (ENSG00000136492) were designed using the E-CRISP Design Tool. Three sgRNAs targeting distinct exons were selected: exon 2 (5′-GGTCTGAATATACAATTGGTG-3′), exon 4 (5′-GTGCCATTTCTTTCAGAAGG-3′), and exon 11 (5′-GCCTCCTCTTTACCATAAAT-3′). As previously described^51^, sequences for Control (CTRL) sgRNAs were taken from Wang et al.^52^: CTRL sgRNA 1 5′-GTGTATCTCAGCACGCTAAC and CTRL sgRNA 2 5′-GAGATTCCGATGTAACGTAC. Complementary oligonucleotides were annealed and cloned into pLentiCRISPR v1 vector (Addgene #49535) according to standard protocols. Correct insertion of sgRNA sequences was verified by Sanger sequencing. Lentiviral particles were produced in HEK293T cells using a second-generation packaging system (psPAX2 and pMD2.G). Viral supernatants were used to transduce HCT116 cells followed by selection with puromycin (1 µg/mL), and clonal cell populations were established by limiting dilution of the transduced and selected cell pools and analyzed for BRIP1 expression by immunoblotting. Genomic DNA (gDNA) from clonal cell lines showing no BRIP1 expression was extracted as previously described^37^. For validation of BRIP1 KO clones, a 783 bp fragment from the BRIP1 locus was amplified by PCR using the following primers: 5′-TTACCCGTCACAGCTTGCTA-3′ and 5′-ACCGACTACCTCAGGATGGA-3′. PCR products were purified using the QIAquick PCR Purification Kit (Qiagen), sequenced or subcloned into the pC2.1 vector (Thermo Fisher Scientific) before sequencing. Sequence analysis was performed by alignment to the *BRIP1* reference sequence to confirm genetic disruption of the BRIP1 locus.

### Generation of HCT116 cells stably expressing HA-BRIP1 and HA-BRIP1^R162Q^

BRIP1 cDNA was derived from pcDNA3-myc-his-BACH1 (Addgene #17642). The patient-derived R162Q variant was generated by site-directed mutagenesis using PCR technology and standard cloning procedures. HA-tagged BRIP1 wild-type (WT) and HA-BRIP1 R162Q lentiviral expression constructs were generated by standard cloning of cDNAs into a modified pLVX-IRES-Puro lentiviral vector (Clontech #632183), and confirmed by Sanger sequencing. Lentiviral particles were produced in HEK293T cells and used for transduction of HCT116 cells and followed by selection with puromycin (2 µg/mL) to establish stable cell pools. Expression of HA-BRIP1 WT and HA-BRIP1 R162Q proteins was validated by immunoblotting using an anti-HA antibody.

### DNA fiber assay

DNA fiber spreading was in principle performed as previously described by Nikolova et al.^53^ including minor modifications. HCT116 cells stably expressing HA-BRIP1 WT or HA-BRIP1 R162Q were seeded in 6-well plates at ∼40–50% confluency one day prior to labeling. Cells were sequentially pulse-labeled with 25 μM 5-chloro-2′-deoxyuridine (CldU; Sigma-Aldrich, Steinheim, Germany) for 25 min followed by 250 μM 5-iodo-2′-deoxyuridine (IdU; TCI Deutschland, Eschborn, Germany) for 25 min. Following labeling, cells were harvested by gentle scraping, and resuspended in 800 μL PBS. Cell suspensions (2 μL) were spotted onto Superfrost™ glass slides and allowed to partially air-dry. DNA fibers were spread by addition of 10 μL spreading buffer (0.5% SDS, 200 mM Tris-HCl pH 7.4, 50 mM EDTA) and incubation at room temperature, followed by tilting of the slides to allow DNA spreading by gravity. Slides were air-dried, fixed in freshly prepared methanol:acetic acid (3:1), air-dried again, and stored at 4 °C. For immunodetection, slides were rehydrated in distilled water and DNA was denatured in 2.5 M HCl for 75 min at room temperature. After washing with PBS, slides were blocked for 1 h at room temperature in 10% (w/v) donkey serum in PBS containing 0.1% Triton X-100. DNA tracts were stained using anti-BrdU antibodies specific for IdU (mouse monoclonal, clone B44, BD Biosciences, #347580) and CldU (rat monoclonal, clone BU1/75 [ICR1], Abcam, #OBT0030), followed by incubation with Alexa Fluor 488 conjugated anti-mouse IgG and Cy3 conjugated anti-rat IgG secondary antibodies (Invitrogen). Slides were washed in PBS and mounted using Mowiol. DNA fibers were visualized using a Zeiss LSM 710 confocal microscope equipped with ZEN 3.8 software. Track lengths were measured using LSM Image Browser (Zeiss), and replication fork speed was calculated using the formula: RS (kb·min⁻¹) = [2.59 × track length (μm)] / pulse time (min). At least 150 replication forks per condition were analyzed from three independent experiments. Replication structures were classified using ImageJ as ongoing forks (CldU–IdU), stalled forks (CldU-only), new origin firing (IdU-only), first-pulse origins (CldU–IdU–CldU), and termination events (IdU–CldU–IdU).

### Immunofluorescence stainings

For immunofluorescence analysis cells were seeded onto glass coverslips. After 24 h, cells were processed as described in the following. γH2AX and phospho-RPA staining: Cells were fixed with 2% paraformaldehyde (PFA) in PBS for 10 min at room temperature, washed with PBS, and permeabilized with 0.1% Triton X-100 in PBS for 10 min. After blocking in 5% goat serum for 1 h at room temperature, coverslips were incubated for 1 h with primary antibodies against γH2AX (Ser139; mouse, 1:400) or phospho-RPA32/RPA2 (Ser8; rabbit, 1:350). After washing, species-specific Cy3-conjugated secondary antibodies (1:400) were applied for 45 min at room temperature in the dark. To-Pro-3 (1:100; Thermo Fisher Scientific) was included for nuclear counterstaining. Samples were washed and mounted in Mowiol. Detection of R-loops (S9.6 staining): For detection of RNA-DNA hybrids, cells were fixed using freshly prepared methanol:acetone (3:1, v/v) for 30 s at room temperature. After blocking in 5% goat serum for 1 h, coverslips were incubated with the S9.6 antibody (mouse monoclonal, Merck, #MABE1095; 1:50 in PBS containing 0.125% Triton X-100) for 2 h at room temperature. Where indicated, recombinant RNaseH1 (10 U per coverslip; New England Biolabs) was added during blocking. After washing, cells were incubated with Cy3-conjugated anti-mouse secondary antibodies (1:500; Invitrogen) together with To-Pro-3 (1:100) for nuclear staining. Coverslips were washed and mounted in Mowiol. Detection of G-quadruplex structures (BG4 staining): G-quadruplex structures were detected by indirect immunofluorescence using the BG4 antibody as described previously^19^. In brief, cells were fixed with 2% PFA for 15 min, permeabilized with 0.7% Triton X-100, and blocked in 0.5% goat serum in PBST. Coverslips were incubated with BG4 antibody (mouse; 1:100) followed by detection using a rabbit anti-FLAG antibody (1:400) and Cy3-conjugated goat anti-rabbit IgG (1:400). To-Pro-3 (1:100) was included for nuclear counterstaining. Samples were washed and mounted in Mowiol. Colocalization of BRIP1 (HA) with γH2AX: For colocalization analysis, HCT116 cells stably expressing HA-BRIP1 WT or HA-BRIP1 R162Q were treated with hydroxyurea (HU) overnight and fixed with 2% PFA. After blocking, cells were incubated with anti-HA (rabbit, polyclonal) and anti-γH2AX (mouse, monoclonal) primary antibodies. Fluorescent detection was performed using Alexa Fluor 488–conjugated anti-rabbit IgG (for HA) and Cy3-conjugated anti-mouse IgG (for γH2AX). Coverslips were imaged by confocal microscopy as indicated.

### Confocal microscopy and mean fluorescence intensity (MFI) measurement

Images were acquired using a Zeiss LSM 710 confocal laser scanning microscope equipped with a Plan-Apochromat 63×/1.4 NA oil immersion objective. All images within each experiment were acquired using identical settings (laser power, detector gain, pinhole size, and scan speed) to allow direct comparison between conditions. Z-stacks were collected using optimal step sizes to ensure complete nuclear coverage. Image analysis and quantification of nuclear foci were performed using ZEN software (Zeiss), applying consistent thresholds across all samples. For each condition, at least 100 nuclei were analyzed from three independent biological replicates. Quantitative data were processed using GraphPad Prism.

The number of foci was quantified manually, or mean fluorescence intensity (MFI) was measured as described^54^. In brief, for MFI analysis, fluorescence images were analyzed using ImageJ. Nuclear regions of interest (ROIs) were manually defined based on To-Pro-3 staining, and MFI was automatically calculated by the software as the mean gray value (average pixel intensity) within each ROI after background subtraction.

### Immunoblotting

Immunoblotting was performed essentially as previously described^55, 56^. In brief, whole-cell lysates were prepared in NP-40/SDS lysis buffer supplemented with protease and phosphatase inhibitors (complete™, β-glycerophosphate, Na₃VO₄, and PMSF). Lysates were sonicated, cleared by centrifugation, and protein concentrations were determined using a BCA assay. Equal amounts of protein were resolved by SDS–PAGE and transferred onto nitrocellulose membranes. Membranes were blocked in 5% BSA or 5% milk and incubated with primary antibodies overnight at 4 °C, followed by incubation with HRP-conjugated secondary antibodies. Signals were detected using enhanced chemiluminescence reagents (Western Lightning Plus-ECL, SuperSignal Dura or Femto) and imaged using an iBright CL1000 system.

### Statistical analysis

All experiments in this study were independently repeated three times (biological replicates). Quantitative data are presented as the mean ± standard deviation. Statistical analyses were performed to evaluate the significance of differences between experimental groups. Since datasets consisted of matched sample pairs measured under different experimental conditions, statistical significance was assessed using a two-tailed paired Student’s T-test. This test was chosen to account for the inherent pairing of samples, minimizing inter-experimental variability and allowing accurate assessment of condition-specific effects. A *p*-value of less than 0.05 was considered statistically significant. Significance levels are represented as follows: *, p < 0.05; **, p < 0.01; ***, p < 0.01; **, p < 0. 001. All statistical analyses and graphical visualizations were carried out using GraphPad Prism software.

## Acknowledgements

We are grateful to Huong Becker for expert technical assistance and Georg Nagel for help with experiments. This work was funded by grants from the Deutsche Forschungsgemeinschaft (DFG) KU3764/3-1 and HO2438/7-1, and the SFB 1361 (Project-ID 393547839), project 19 (to TGH).

## Author contributions

S.K, T.N., R.A, M.K., P.-O.F., and T.G.H., designed the experiments. S.K., T.N., D.P., S.N., and P.-O.F. performed the experiments. S.K., T.N., S.N., M.K., and P.-O.F., analyzed the data. S.K., and T.G.H., wrote the manuscript. S.K., and T.G.H.., supervised the research.

