## Supplementary Figures for "Role of a childhood cancer-linked BRIP1/FANCJ germline variant in genomic instability and cancer cell vulnerability"

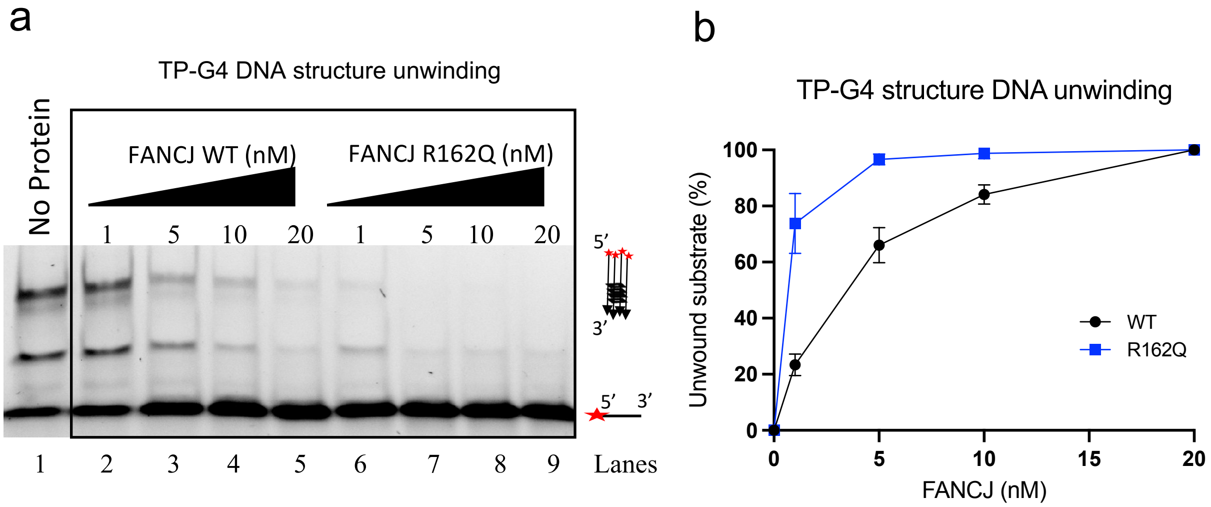


**Suppl. Fig. 1**. **BRIP1/FANCJ R162Q is hyper-active in G4 unwinding. a**, Representative native 6% polyacrylamide gel of DNA unwinding assay with indicated concentrations of FANCJ WT and R162Q variant with TP-G4 DNA structures. The *indicates the position of FAM labelling at the 5’-end of the TP-G4 oligo (49-mer). **d**, Quantitation of experiments such as shown in a. n=3 independent experiments. Data are mean +/– SEM.

Karbassi et al., Supplementary Figure 1

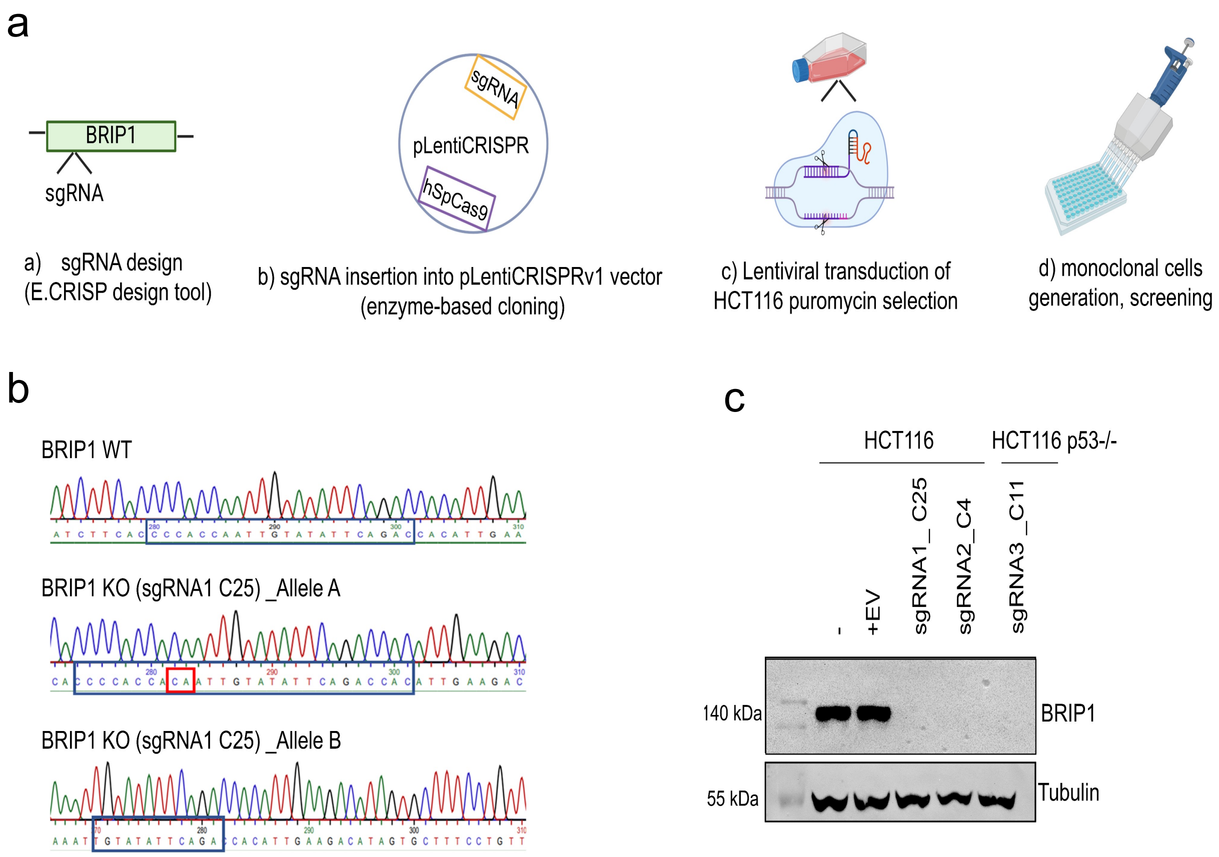


Karbassi et al., Supplementary Figure 2
